# DeepStrain: A Deep Learning Workflow for the Automated Characterization of Cardiac Mechanics

**DOI:** 10.1101/2021.01.05.425266

**Authors:** Manuel A. Morales, Maaike van den Boomen, Christopher Nguyen, Jayashree Kalpathy-Cramer, Bruce R. Rosen, Collin M. Stultz, David Izquierdo-Garcia, Ciprian Catana

## Abstract

Myocardial strain analysis from cinematic magnetic resonance imaging (cine-MRI) data could provide a more thorough characterization of cardiac mechanics than volumetric parameters such as left-ventricular ejection fraction, but sources of variation including segmentation and motion estimation have limited its wide clinical use. We designed and validated a deep learning (DL) workflow to generate both volumetric parameters and strain measures from cine-MRI data, including strain rate (SR) and regional strain polar maps, consisting of segmentation and motion estimation convolutional neural networks developed and trained using healthy and cardiovascular disease (CVD) subjects (n=150). DL-based volumetric parameters were correlated (>0.98) and without significant bias relative to parameters derived from manual segmentations in 50 healthy and CVD subjects. Compared to landmarks manually-tracked on tagging-MRI images from 15 healthy subjects, landmark deformation using DL-based motion estimates from paired cine-MRI data resulted in an end-point-error of 2.9 ± 1.5 mm. Measures of end-systolic global strain from these cine-MRI data showed no significant biases relative to a tagging-MRI reference method. On 4 healthy subjects, intraclass correlation coefficient for intra-scanner repeatability was excellent (>0.95) for strain, moderate to excellent for SR (0.690-0.963), and good to excellent (0.826-0.994) in most polar map segments. Absolute relative change was within ~5% for strain, within ~10% for SR, and <1% in half of polar map segments. In conclusion, we developed and evaluated a DL-based, end-to-end fully-automatic workflow for global and regional myocardial strain analysis to quantitatively characterize cardiac mechanics of healthy and CVD subjects based on ubiquitously acquired cine-MRI data.

## I. INTRODUCTION

CARDIAC mechanics reflects the precise interplay between myocardial architecture and loading conditions that is essential for sustaining the blood pumping function of the heart. The ejection fraction (EF) is often used as a left-ventricular (LV) functional index, but its value is limited when mechanical impairment occurs without an EF reduction [1]. Alternatively, tissue tracking approaches for strain analysis provide a more thorough characterization through non-invasive evaluation of myocardial deformation from echocardiography or cinematic magnetic resonance imaging (cine-MRI) data [2], and could be used to identify dysfunction before EF is reduced [3]. Unfortunately, various sources of discrepancies have limited the wide clinical applicability of these techniques, including factors related to imaging modality, algorithm, and operator [4]. More accurate measures could be obtained from tagging-MRI data widely regarded as the reference standard for strain quantification [5], [6], but collection of these data requires highly specialized and complex sequences that have mainly remained research tools, whereas echocardiography and cine-MRI data are ubiquitously acquired in clinical practice.

Irrespective of algorithm or modality, e.g., speckle tracking for echocardiography or feature tracking for cine-MRI, the main challenge is to estimate motion within regions along the myocardial wall [2]. Operator-related discrepancies are introduced when the myocardial wall borders are delineated manually, a time-consuming process that requires considerable expertise and results in significant inter- and intra-observer variability [7], [8]. Automatic delineation approaches have been implemented within computational pipelines [9], but other factors related to motion tracking algorithms also influence strain assessment, including the appropriate selection of tuneable parameters whose optimal values can differ between patient cohorts and acquisition protocols (e.g., the size of the search region in block-matching methods [10]). Further, these algorithms often make assumptions about the properties of the myocardial tissue (e.g., incompressible and elastic [11], [12]), or use registration methods to drive the solution towards an expected geometry. However, recent evidence has shown the validity of these assumptions varies between healthy and diseased myocardium [13], [14], suggesting these approaches may not accurately reflect the underlying biomechanical motion [14]. Lastly, modality-related image quality could complicate interpretation of abnormal strain values since these could reflect either real dysfunction or artifact-related inaccuracies, leading to some degree of subjectivity or non-conclusive results [3].

Deep Learning (DL) methods have demonstrated the advantage of allowing real-world data guide learning of abstract representations that can be used to accomplish pre-specified tasks, and have been shown to be more robust to image artifacts than non-learning techniques for some applications [15], [16]. DL segmentation methods have been proposed [17]–[20] and implemented within strain computational pipelines [21], [22], and recent studies have shown that cardiac motion estimation can also be recast as a learnable problem [23]–[26]. These methods usually consist of an intensity-based loss function and a constrain term [23], [27], the latter using common machine learning techniques (e.g., L2 regularization of all learnable parameters [24]) or direct regularization of the motion estimates (e.g., smoothness penalty [23], anatomy-aware [26]). However, because ground-truth cardiac motion is challenging to acquire, whether these constrains improve the accuracy of motion or strain estimates is not yet clear. Further, the added-value of DL-based regional strain estimation has not been demonstrated.

We have recently developed a learning method for cardiac motion estimation that produces more accurate estimates than various techniques, including B-spline, diffeomorphic, and mass-preserving algorithms [28], and showed these estimates could potentially be used to detect regional dysfunction. Thus, incorporating our method within a strain analysis framework could potentially enable accurate, user-independent, and quantitative characterization of cardiac mechanics at a both global and regional level. Once trained, such method would not necessitate further parameter tunning or optimization, which is time-consuming for larger datasets and daily clinical practice. While this framework could be based on echocardiography images [29], these data remain limited for strain mapping tasks by their low reproducibility of acquisition planes [4] and temporal stability of tracking patterns [30]. In contrast, cine-MRI offers the most accurate and reproducible assessment of cardiac anatomy and function, thus providing a more thorough set of data for learning-based motion models.

We propose DeepStrain, an automated workflow that derives global and regional strain measures from cine-MRI data by decoupling motion estimation and segmentation tasks. After verifying the effects of smoothing and anatomical regularizers on motion and strain, convolutional neural networks for pre-processing (i.e., centering and cropping), segmentation, and motion estimation were implemented, trained, validated, and compared to state-of-the-art methods. Finally, accuracy of strain values was assessed using a tagging-MRI algorithm as reference standard, intra-scanner repeatability was measured using subjects with repeated scans, and potential clinical applications of global and regionals myocardial strain measures were demonstrated on patient populations.

## II. Method

### A. Datasets

For development we used the *Automated Cardiac Diagnosis Challenge (ACDC)* dataset [31], consisting of cine-MRI data from 150 subjects evenly divided into five groups: healthy and patients with hypertrophic cardiomyopathy (HCM), abnormal right ventricle (ARV), myocardial infarction with reduced ejection fraction (MI), and dilated cardiomyopathy (DCM). These data were publicly available as train (*n*=100) and test (*n*=50) sets, with manual segmentations included for the train set only. For validation of motion and strain measures we used the *Cardiac Motion Analysis Challenge (CMAC)* dataset [32], consisting of paired tagging- and cine-MRI data from 15 healthy subjects. To assess intra-scanner repeatability, four healthy volunteers were recruited to undergo repeated scans on a 3T MRI scanner. All cine-MRI frames and corresponding segmentations were resampled to a 256×256×16 volume grid with 1.25 mm × 1.25 mm in-plane resolution and variable slice thickness (4-7 mm). See supplementary section S1 for acquisition protocol.

### B. Myocardial Strain Definitions

Strain represents percent change in myocardial length per unit length. The three-dimensional (3D) analog for MRI is given by the Lagrange strain tensor

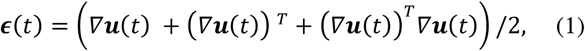

where ***u*** (*t*) denotes myocardial displacement from a fully-relaxed end-diastolic phase at *t=0,* to a contracted frame at *t>0.* Radial and circumferential strain are the diagonal components of the tensor ***ϵ*** evaluated in cylindrical coordinates. *Strain rate (SR)* is the time derivative of (1). *Global strain* is defined as the average of ***ϵ*** over the whole LV myocardium (LVM) volume. *Regional* strain is defined as the average of ***ϵ*** over the volume of specific LVM segments defined by the American Heart Association (AHA) polar map [33], which requires labels of the right ventricle to construct. Specific parameters based on timing and magnitude are extracted from the measures evaluated over a whole cardiac cycle: *end-systolic strain (ESS)*, defined as the global strain value at end-systole; *systolic strain rate (SRs)*, defined as the peak (i.e., maximum) absolute value of global SR during systole; *early-diastolic strain rate (SRe)*, defined as the peak absolute value of global SR during diastole.

### C. Centering, Segmentation, and Motion Estimation

DeepStrain (Fig. 1) consists of a series of convolutional neural networks that perform three tasks: a ventricular centering network (VCN) for automated centering and cropping, a cardiac motion estimation network (CarMEN) to generate ***u***, and a cardiac segmentation network (CarSON) to generates tissue labels. Estimates of ***u*** are used to calculate myocardial strain, and segmentations are used to derive volumetric parameters, identify a cardiac coordinate system for strain analysis, and generate tissue labels used for anatomical regularization of the motion estimates at training time.

**Fig. 1.**
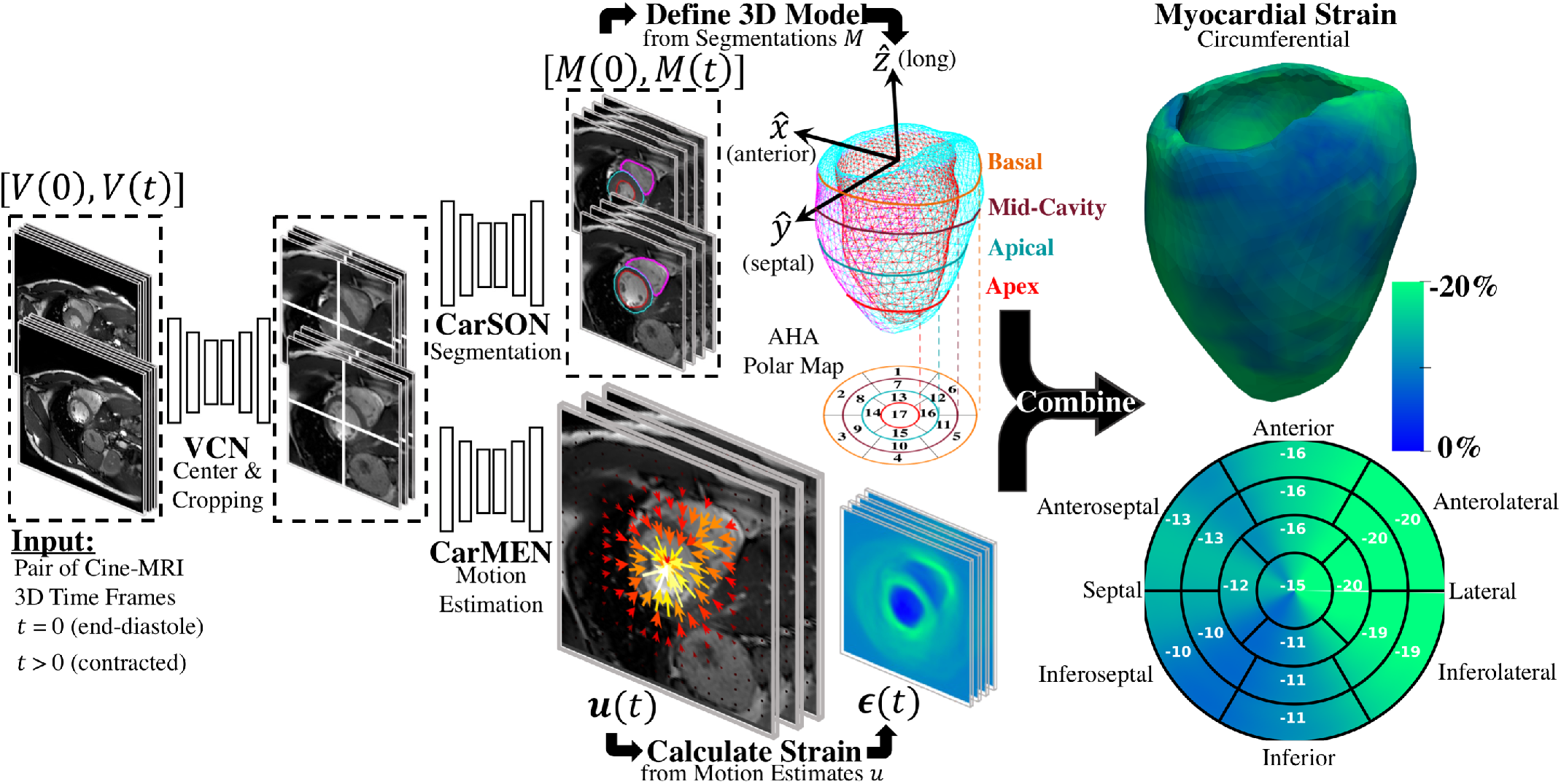
Overview of proposed DeepStrain workflow. VCN centers and crops the input pair of cine-MRI frames. Tissue labels generated by CarSON are used to build an anatomical model. Motion estimates derived from CarMEN are used to calculate strain measures, and these estimates are combined with the anatomical model to enable global and regional strain analyses.

Let *V_t_* be a cine-MRI frame at time *t* defined over a 3D spatial domain *Ω* ⊂ ℝ^3^. Using a pair of frames *V*_0_, *V_t_* as an input, VCN centers and crops the images around the center of mass of the LV, CarSON generates segmentations *M*_0_, *M_t_* of the LV, RV, and LVM, and CarMEN estimates the motion ***u**_t_* of the heart from *V*_0_ to *V_t_*. Thus, for each voxel *p* ∈ *Ω*, ***u**_t_* (*p*) is an approximation of the myocardial displacement during contraction such that *V*_0_(*p*) and (***u**_t_* ∘ *V_t_*)(*p*) correspond to similar cardiac regions. The operator ∘ refers to application of a spatial transform to *V_t_* using ***u**_t_* via trilinear interpolation [34].

#### 1) Architectures

All networks have a common encoder-decoder architecture consisting primarily of convolution, batch normalization [35], and PReLU [36] layers with residual connections [37] (see supplementary section S2). Briefly, VCN is a 3D architecture that uses a single-channel array *V* with size 256×256×16 to generate a single-channel array *G_pred_* of equal size, where *G_pred_* corresponds to a Gaussian distribution with mean defined as the LVM center of mass. V is centered and cropped around the voxel with the highest value in *G_pred_* to generate a new cropped array of size 128×128×16, which is then the input to segmentation and motion estimation networks. CarSON is a two-dimensional (2D) architecture that uses images of size 128×128 to generate a 4-channel segmentation *M_pred_* of equal size, each channel corresponding to a label. CarMEN uses a 2-channel input volume, consisting of two concatenated arrays with size 128×128×16, to generate a 3-chanel array ***u*** of equal size. Each channel in ***u*** represents the *x*, *y* and *z* components of motion.

#### 2) Loss Functions

VCN was evaluated using the mean square error

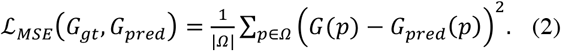

For CarSON, we implemented a multi-class Dice coefficient function

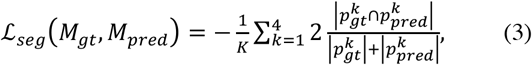

where *k* ∈ [1, 4] represents each of the tissue labels, and *p*^*k*^ ∈ *M* denotes all the pixels with label *k*. A combination of three functions was used for motion estimation. First, we used an unsupervised loss function 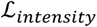 that evaluates CarMEN using the input volumes and generated motion estimates

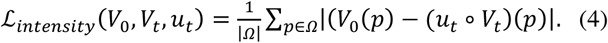

Second, we used a supervised function 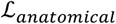 that leverages segmentations of the input volumes at training time to impose an anatomical constrain on the estimates

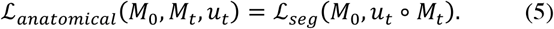

Third, smooth estimates were encouraged by using a diffusion regularizer

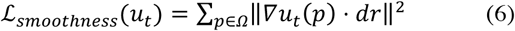

where *dr* is the spatial resolution of *V* and accounts for differences between in-plane and slice resolution. Thus, the loss function for CarMEN is a linear combination of (4), (5), and (6), weighted by *λ_i_*, *λ_a_*, *λ_s_*, accordingly.

We conducted optimization experiments using synthetic data [38], [39] to assess the impact of smoothing and anatomical regularization on motion and strain estimates (supplementary section S3). These experiments showed smoothness improves the accuracy of the motion vectors *direction*, and anatomical regularization improves the *magnitude* of the vectors relative to the ground-truth motion (see supplementary Fig. S1 and S2). The optimal values *λ*_*i*_ = 0.01, *λ*_*a*_ = 0.5, *λ*_*s*_ = 0.1 were used to train CarMEN.

#### 3) Training and Testing

Networks were trained in TensorFlow ver. 2.0 with Adam optimizer parameters beta1, 2 = 0.9,0.999, batchsize = 80 (5 for CarMEN), and epochs = 1000 (300 for CarMEN). Ground-truth distributions for VCN were created using the manual segmentations. VCN and CarSON were trained using the end-diastolic and end-systolic frames of the train set, as only these included ground-truth segmentations. This provided 200 training samples for VCN and 3200 for CarSON, the latter having more samples since it is a 2D architecture and all frames were resampled to a volume with 16 slices. VCN was tested by five-fold cross-validation, whereas the accuracy of CarSON was assessed by submitting the results to the challenge website.

Once CarSON was trained, we generated segmentations of the test set to train CarMEN using the entire ACDC dataset. Only the [end-diastolic, end-diastolic] and [end-diastolic, end-systolic] pairs were used. The former is essential for the network to adequately learn how to scale the motion vectors, i.e., motion should be exactly zero if the frames are equal. The entire cycle is analyzed at testing time by using sequential input pairs [*V*_0_, •] that kept the end-diastolic frame constant while we varied *V_t_* for all time frames *t* > 0. Using this approach ***u**_t_* was derived for all times. Data augmentation included random rotations and translations, random mirroring along the x and y axes, and gamma contrast correction. All data augmentation was performed only in the x-y plane.

### D. Evaluation

#### 1) Segmentation and Motion Estimation

CarSON and manual segmentations were compared using the Hausdorff distance (HD) and Dice Similarity Coefficient (DSC) metrics at both end-diastole and end-systole. Accuracy of LV volumetric measures derived from segmentations, including end-diastolic volume (EDV), EF, and LVM, was assessed using the correlation, bias, and standard deviation metrics. The mean absolute error (MAE) for the LV EDV and LVM were also computed for comparison against the intra- and inter-observer variability reported by [31]. We compared our results to top-3 ranked methods published for the ACDC test set as these appear in the leader-board of the challenge [17]–[20].

The CMAC organizers defined 12 landmarks at the intersection of gridded tagging lines at end-diastole on tagging images, one landmark *p*_0_ per wall per ventricular level. These landmarks were manually-tracked by two observers over the cardiac cycle. Conversion from tagging to cine coordinates was done using DICOM header information. We used the CarMEN motion estimates *u_t_* to automatically deform the landmarks at end-diastole, and the accuracy was assessed using the in-plane end-point error (EPE) between deformed 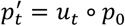 and manually-tracked *p_t_* landmarks, defined by

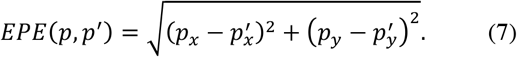

Due to temporal misalignment between the tagging and cine acquisitions, EPE was evaluated only at end-systole (*t* = *t_ES_*). Specifically, let *p_ij_* (*t*) denote the manually-tracked landmarks of subject *i* at frame *t* by observer *j*. The accuracy of CarMEN was assessed using the average EPE

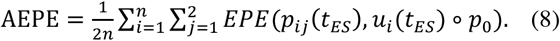

Our results were compared to those reported by the four groups that responded to the challenge [32], MEVIS [40], IUCL [9], UPF [11], and INRIA [12], [41]. All groups submitted tagging-based motion estimates, but only UPF and INRIA provided estimates based on cine-MRI.

#### 2) Strain Validation and Intra-Scanner Repeatability

The tagging-MRI method with the lowest AEPE was used as the reference for strain analysis. The tagging-MRI-based motion estimates were registered and resampled to the cine-MRI space. Global strain and SR values throughout the entire cardiac cycle were derived from the resampled estimates as described in [42].

Global- and regional-based analyses were performed to assess the repeatability of measures from two acquisitions. Relative changes (RC) and absolute relative changes (aRC) were calculated, taking the first acquisition as the reference. ESS and SR were calculated for the global-based analysis, and for region-based analyses, ESS values were normalized using the AHA polar map, and both RC and aRC were evaluated for each of the segments in the polar map.

#### 3) Statistics

Bland-Altman analysis was used to quantify agreement between predicted and tagging strain measures. We used the term *bias* to denote the mean difference and the term *precision* to denote the standard deviation of the differences. Differences were also assessed using a paired *t*-test with Bonferroni correction for multiple comparisons. For global- and regional-based analyses of intra-scanner repeatability, ICC estimates and their 95% confidence intervals (CI) were calculated based on a single-rating, absolute agreement, 2-way mixed-effects model. Analyses were performed on python v3.4 [43].

## III. Results

### A. Segmentation and Motion Estimation

Centering, segmentation, and motion estimation for an entire cardiac cycle (~25 frames) was accomplished in <13s on a 12GB GPU and <2.2 min on a 32GB RAM CPU. VCN located the LV center of mass with a median error of 1.3 mm.

Correlation of CarSON and manual LV volumetric measures was >0.98 across all measures (Table 1), and biases in EF (+0.25 ± 3.2%), end-diastolic (+0.76 ± 6.7 mL) and end-systolic (+0.19 ± 5.8 mL) volumes, and mass (+1.4 ± 10.3 g) were not significant. Further, these biases were smaller than those obtained with other methods, which were positive for LV EDV (1.5 to 3.7 mL), negative for LVM (−2.1 to −2.9 g), and close to zero (±0.5%) for EF. Simantiris *et al*. [17] obtained the best precision for LV EF (2.7 vs. 3.2% variance with CarSON), EDV (4.6 vs. 6.7 mm), and LVM (6.5 vs. 10.3 g). Isensee *et al*. [18] obtained the best results on geometric metrics, i.e., lower HD for the LV (end-diastole 5.5 vs. 5.7 mm; end-systole 6.9 vs. 7.7 mm) and LVM (7.0 vs. 8.1 mm; 7.3 vs. 9.2 mm), and higher DSC for the LVM (0.904 vs. 0.898; 0.923 vs. 0.913). The DSC for the LV was similar for all methods (~0.967, ~0.929). MAE for the LV EDV and LVM were 5.3 ± 4.1 mL and 6.8 ± 6.5 g.

**Table I.** **State-Of-The-Art Methods** For Left-Ventricular Segmentation Shown At End-Diastole (ED) And End-Systole (ES) On The ACDC Test Set Compared To Proposed Approach. **Red** Are The Best Results For Each Metric.

| Left-Ventricle Label |  | Dice Similarity Coefficient |  | Hausdorff Distance |  | Ejection Fraction |  |  | End-Diastolic Volume |  |  |
| --- | --- | --- | --- | --- | --- | --- | --- | --- | --- | --- | --- |
| Rank | Method | ED val. | ES val. | ED mm | ES mm | Corr. val. | bias±std % | % | Corr. val. | bias±std mL | mL |
| * | **CarSON** | **0.967** | **0.929** | 5.656 | 7.676 | 0.990 | **0.252** | 3.183 | 0.996 | **0.762** | 6.672 |
| 1 | Simantiris *et al.* [17] | **0.967** | 0.928 | 6.366 | 7.573 | **0.993** | -0.360 | **2.689** | **0.998** | 2.032 | **4.611** |
| 2 | Isensee *et al.* [18] | **0.967** | 0.928 | **5.476** | **6.921** | 0.991 | 0.490 | 2.965 | 0.997 | 1.530 | 5.736 |
| 3 | Zotti *et al.* [19] | 0.964 | 0.912 | 6.180 | 8.386 | 0.990 | -0.476 | 3.114 | 0.997 | 3.746 | 5.146 |
| Myocardium Label |  | Dice Similarity Coefficient |  | Hausdorff Distance |  | Left-Ventricular Mass |  |  | End-Systolic Volume |  |  |
| Rank | Method | ED val. | ES val. | ED mm | ES mm | Corr. val. | bias±std g | g | Corr. val. | bias±std mL | mL |
| * | **CarSON** | 0.898 | 0.913 | 8.128 | 9.189 | 0.981 | **1.405** | 10.32 | 0.985 | **1.152** | 9.391 |
| 1 | Isensee *et al.* [17] | **0.904** | **0.923** | **7.014** | **7.328** | 0.987 | -2.547 | 8.28 | 0.988 | -1.984 | **8.335** |
| 2 | Simantiris *et al.* [18] | 0.891 | 0.904 | 8.264 | 9.575 | **0.992** | -2.904 | **6.46** | 0.983 | -2.134 | 10.11 |
| 3 | Calisto *et al.* [20] | 0.873 | 0.895 | 8.197 | 8.318 | 0.989 | -2.1 | 7.91 | **0.988** | -1.79 | 8.575 |

Fig. 2a illustrates a representative example of the tagging and cine images from a CMAC subject. Landmarks defined at end-diastole were deformed to end-systole using the CarMEN estimates and compared to manual tracking. Banding artifacts on cine images showed no clear effect on derived motion estimates or landmark deformation, as shown in end-systole (Fig. 2a, yellow arrow) or throughout the whole cardiac cycle (see supplementary video). The manual tracking inter-observer variability was 0.86 mm (Fig. 2b, dotted line). Within cine-based techniques, CarMEN (2.89 ± 1.52 mm) and UPF (2.94 ± 1.64 mm) had lower (p<0.001) AEPE relative to INRIA (3.78 ± 2.08 mm), but there was no significant difference between CarMEN and UPF. All tagging-based methods had lower AEPE compared to cine approaches, particularly MEVIS (1.58 ± 1.45 mm) and UPF (1.65 ± 1.45 mm).

**Fig. 2.**
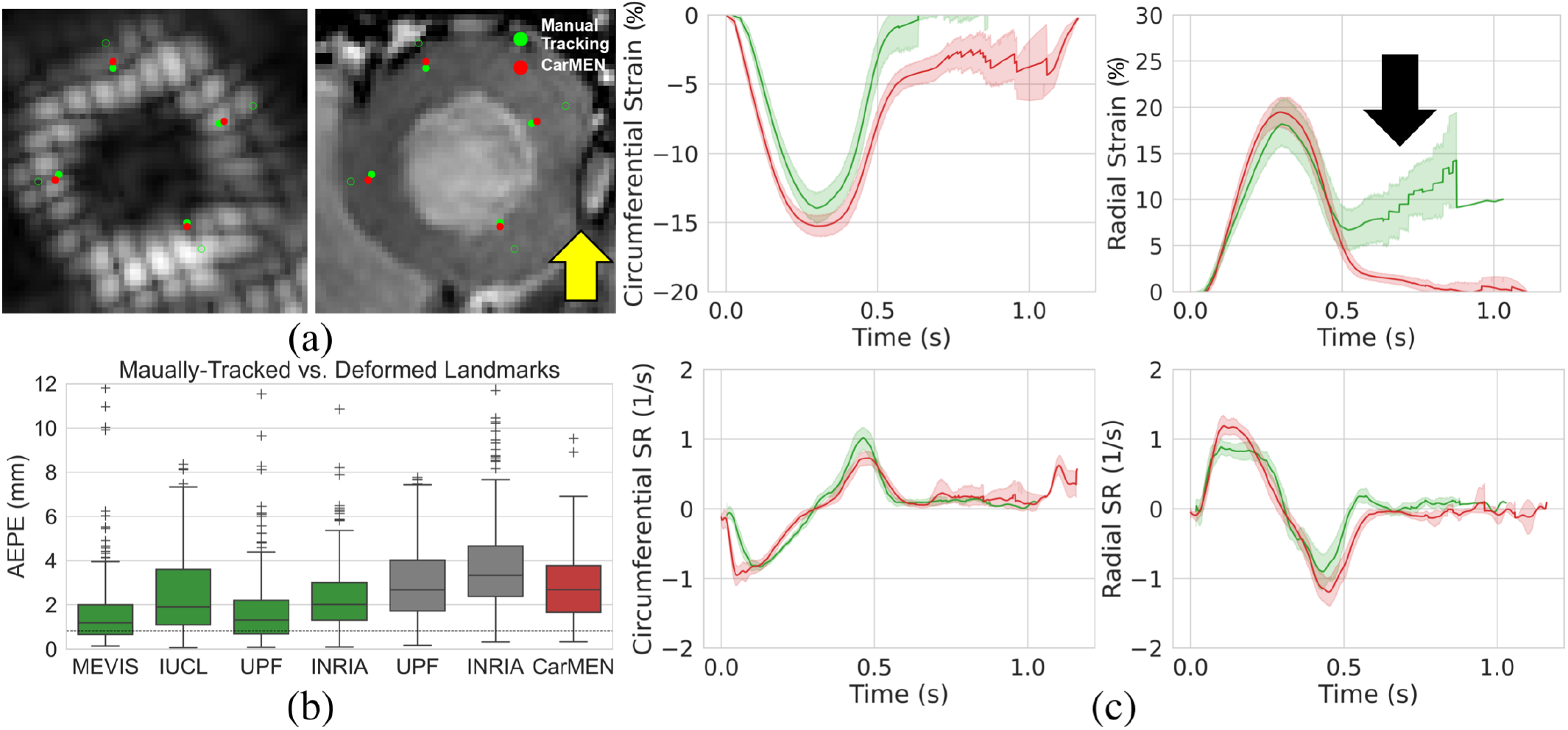
Validation of motion and strain. (a) Landmarks at end-diastole (unfilled green) are manually-tracked (green) and deformed with CarMEN to end-systole (red). Yellow arrow indicates a banding artifact. (b) Average end-point-error (AEPE) was assessed and compared to other methods. (c) MEVIS- and DeepStrain-based strain (top) and strain rate (SR, bottom) measures are compared. Black arrow shows strain inaccuracies with MEVIS.

### B. Strain Analysis

Table 2 shows the normal ranges (mean [95% CI]) of strain derived from cine-MRI data for all healthy subjects, including subjects from the training, validation, and repeatability cohorts. DeepStrain generated values with narrow CI for circumferential (~1%) and radial (~2%) ESS, and circumferential (~0.15 s^−1^) and radial (~0.25 s^−1^) SR. Specifically, circumferential and radial values across datasets were: −16.9% [−17.6 −16.3] and 22.6% [21.4 23.8] for ESS, −1.12 s^−1^ [−1.19 −1.05] and 1.30 s^−1^[1.20 1.40] for SRs, and 0.76 s^−1^ [0.69 0.83] and −1.38 s^−1^ [−1.51 −1.24] for SRe, accordingly. These values were similar to those from tagging-based ones, although circumferential SRe from cine-MRI data was lower, mostly in the train set (0.7± 0.2 s^−1^).

**Table II.**
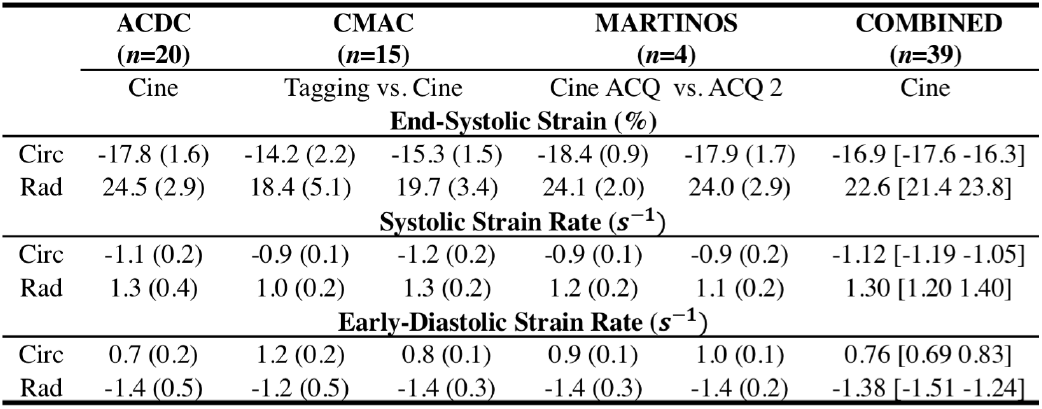
**Normal Ranges Of Strain** With DeepStrain In Healthy Subjects. Tagging-Based Measures Are Shown For The CMAC Cohort. DeepStrain Repeatability Is Shown For Two Acquisitions (ACQ).

Comparison of tagging- and cine-based strain measures with matched subjects showed an overall agreement in timing and magnitude of strain and SR throughout the cardiac cycle, although tagging-based measures of radial ESS diverge after early diastole (Fig. 2c, black arrow), and there were visual differences in peak SR parameters. Visual inspection of image artifacts on cine data showed no clear evidence that these artifacts affected strain values derived with DeepStrain (see supplementary Fig. S3). Quantitative comparisons of tagging- and cine-based measures showed biases in circumferential ESS (−14.2 ± 2.2 vs. −15.3 ± 1.5%; bias −1.17 ± 2.93%), radial ESS (18.4 ± 5.1 vs. 19.7 ± 3.4%; +1.26 ± 5.37%) and SRe (−1.2 ± 0.5 vs. −1.4 ± 0.3; −0.21 ± 0.52 s^−1^) were small and not significantly different from zero (see supplementary Fig. S4). However, there were larger differences (p<0.01) in radial SRs (1.0 ± 0.2 vs. 1.3 ± 0.2 s^−1^; 0.32 ± 0.34 s^−1^), and circumferential SRs (−0.9 ± 0.1 vs. −1.2 ± 0.2 s^−1^; 0.30 ± 0.22 s^−1^) and SRe (1.2 ± 0.2 vs. 0.8 ± 0.1 s^−1^; 0.40 ± 0.23 s^−1^).

Representative strain measures of a single subject derived from two acquisitions are shown in Fig. 3. The AHA polar maps from both acquisitions showed comparable regional variations in ESS, particularly for circumferential ESS in the inferoseptal wall (Fig. 3a, orange arrows). Global curves throughout the entire cardiac cycle also showed visual agreement in both timing and magnitude (Fig. 3b). From these data, circumferential (−14.1 vs. −14.3%) and radial (17.9 vs. 17%) ESS (Fig. 3b, purple), circumferential SRs (0.95 vs. 0.90 s^−1^) and SRe (−0.74 vs. −0.82 s^−1^), and radial SRs (1.03 vs. 1.08 s^−1^) and SRe (−1.12 vs. −1.11 s^−1^) global parameters were also found to be similar (Fig. 3b, yellow). In addition, while not quantified in this study, the late-diastolic filling peaks were also comparable (Fig. 3b, blue).

**Fig. 3.**
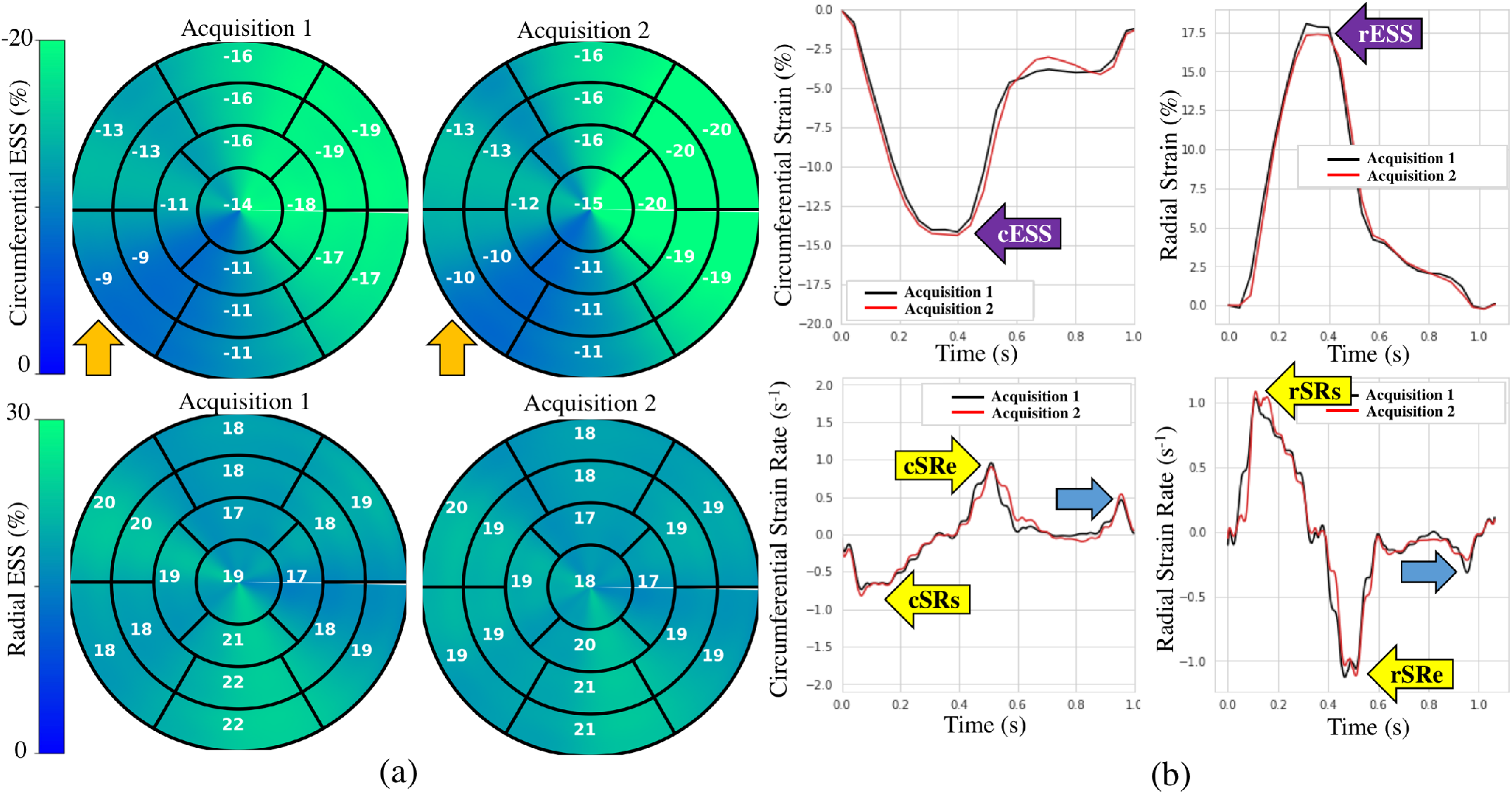
Global and regional strain measures of representative subject. (a) Regional end-systolic strain measures show visual agreement (orange arrow). (c) Global strain and strain rate (SR) measures also show visual agreement.

Table 3 shows the RC, aRC, ICC, and LoA across subjects for the global parameters. The average aRC was below 5% for ESS (circumferential: 3.1 ± 1.8%; radial: 4.3 ± 3.4%), below 7% for SRs (5.7 ± 4.4%; 6.9 ± 10.4%), and below 11% for SRe parameters (10.2 ± 8.8%; 3.8 ± 3.1%). ICC results showed repeatability was excellent for ESS (0.954; 0.968), good for SRs (0.889; 0.754), moderate for circumferential SRe (0.690), and excellent for radial SRe (0.963) values. The LoA, which defines the interval where to find the expected differences in 95% of the cases assuming normally distributed data, were ~1% and ~4% for circumferential and radial ESS, and <0.05 s^−1^ for all SR measures.

**Table III.** **Intra-Scanner Repeatability** Of Global Circumferential (Circ) And Radial (Rad) Strain Measures.

| Measure | RC (%) | aRC (%) | ICC [95% CI] | LoA |
| --- | --- | --- | --- | --- |
| circ ESS | 3.13 (1.81) | 3.13 (1.81) | 0.954 [0.470-1.000] | [0.0 1.0 %] |
| rad ESS | 2.34 (5.32) | 4.27 (3.37) | 0.968 [0.590-1.000] | [-2.1 1.3 %] |
| circ SRs | -2.45 (7.33) | 5.66 (4.37) | 0.889 [0.050-0.990] | [-0.010 0.013 s<sup>-1</sup>] |
| rad SRs | -4.33 (12.12) | 6.87 (10.44) | 0.754 [0.370-0.980] | [-0.019 0.030 s<sup>-1</sup>] |
| circ SRe | 8.98 (10.48) | 10.22 (8.84) | 0.690 [0.480-0.980] | [-0.009 0.024 s<sup>-1</sup>] |
| rad SRe | -2.73 (4.36) | 3.83 (3.07) | 0.963 [0.540-1.000] | [-0.016 0.007 s<sup>-1</sup>] |

The ESS, RC, and aRC maps averaged across subjects are shown in Fig. 4. Visually, these maps (Fig. 4b) showed the average RC and aRC were marginal (~1%) in more than half of the polar map segments. Specifically, values were marginal for circumferential ESS (~1%) in the anterior, anteroseptal, and anterolateral walls, but were larger in the inferior region, most notably in the basal- and mid-inferoseptal segments (7%). For radial ESS the largest changes were found in the mid-anterolateral segment (6%), whereas changes in the anteroseptal, inferior and inferolateral walls were very small (~1%). The RC and aRC per subject are provided in boxplot form in supplementary Fig S5. These results showed that, in most of the segments, the RC and aRC were less than ~10%, although larger differences were noted in the inferoseptal wall for radial ESS, and anterolateral wall for circumferential. Supplementary Table S1 shows the ICC and LoA per segment, including the whole-map average. For radial ESS, the ICC results showed excellent repeatability across all segments. Circumferentially, all segments showed good to excellent repeatability, except for the basal- and mid-inferolateral segments. LoAs showed that 95% of differences occurred within ~3% and ~4% intervals for circumferential and radial ESS.

**Fig. 4.**
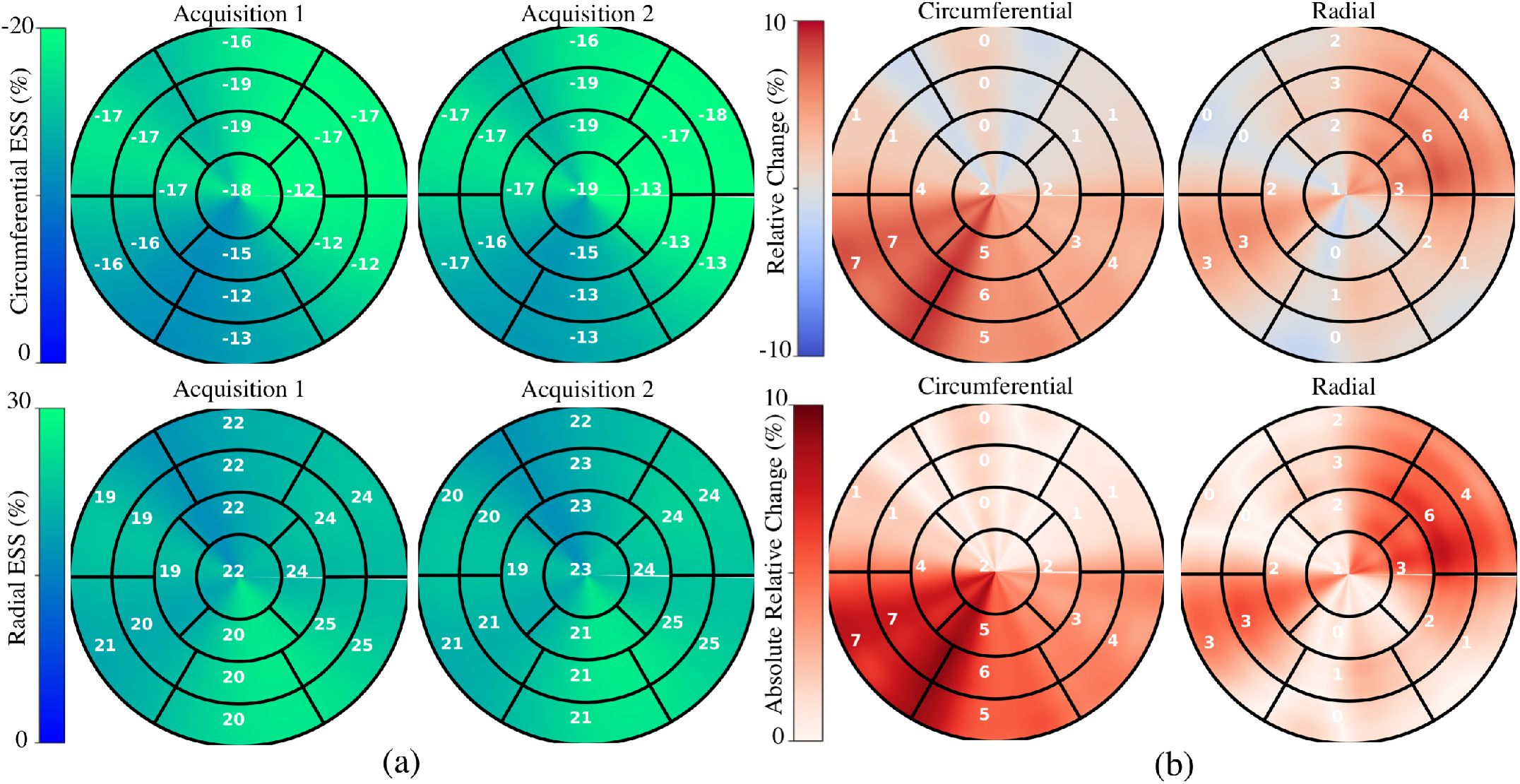
Intra-scanner repeatability of regional myocardial strain measures. (a) Average of subject-specific regional end-systolic strain (ESS) maps during two acquisitions. (b) Average changes between acquisitions.

### C. Evaluation in Patients with Cardiovascular Disease

Regional measures of ESS averaged over patient population (see supplementary figure S6), as well as global values of strain and SR across the cardiac cycle (Fig. 5) for all 100 subjects in the ACDC train set showed progressive decline in strain values starting with HCM, followed by ARV, MI, and DCM. Specifically, relative to the healthy group, radial ESS was reduced in all patient populations. Radial systolic and early-diastolic SR were also reduced in all patient groups, except for systolic SR in HCM. Fig. 6 shows both the cine-MRI image and the circumferential ESS polar map of a healthy subject and two patients with MI. Strain values in the healthy polar map have a homogeneous distribution. In contrast, in one MI patient the map indicates a diffused reduction, and inspection of the myocardium on the cine-MRI image shows an anteroseptal infarct that coincides in location with segments with more prominent decreases in strain. In a different MI patient with an infarct located in a similar septal region, strain changes are focal and localized to the anteroseptal wall.

**Fig. 5.**
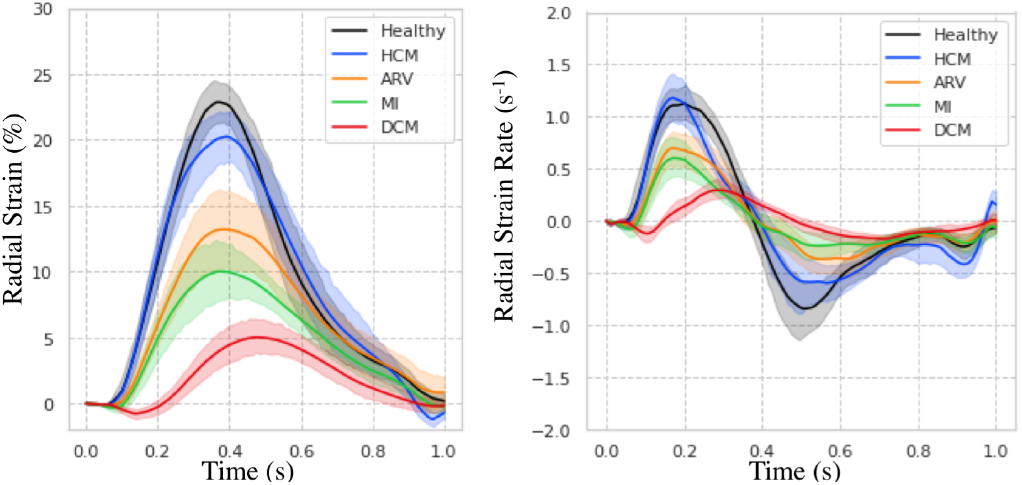
Strain and strain rate measures computed on the ACDC train set.

**Fig. 6.**
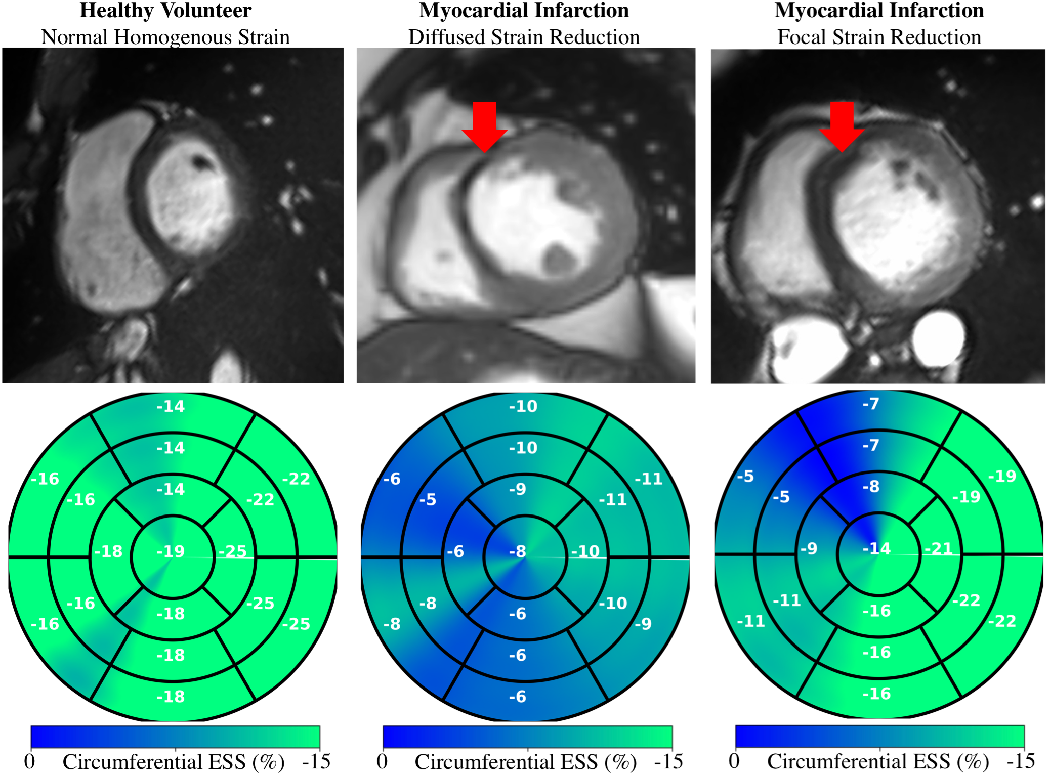
Regional strain in healthy and patients with MI. Myocardial infarction can result in diffused (center) and focal (right) strain reduction.

## IV. DISCUSSION

Learning-based methodologies have the potential to meet the technical challenges associated with myocardial strain analysis. In this study we developed a fast DL framework for strain analysis based on cine-MRI data that does not make assumptions about the underlying physiology, and we benchmarked its segmentation, motion, and strain estimation components against the state-of-the-art. We compared our segmentations to other DL methods, motion estimates to other non-learning techniques, and strain measures to a reference tagging-MRI technique. We also presented the intra-scanner repeatability of DeepStrain-based global and regional strain measures, and showed that these measures were robust to image artifacts in some cases. Global and regional applications were also presented to demonstrate the potential clinical utilization of our approach.

### A. Volumetric Measures

Segmentation from MRI data is a task particularly well suited for convolutional networks given the excellent soft-tissue contrast, thus all top performing methods on the ACDC test set were based on DL approaches (Table 1). Isensee *et al*. [18] had remarkable success on geometric metrics, but this and other approaches result in a systematic overestimation of the LV EDV and thus underestimation of LVM. In contrast, CarSON generated less biased measures of LV volumes and mass, which were not significant. Although Simantiris *et al*. [17] obtained the most precise measures, possibly due to their extensive use of augmentation using image intensity transformations, across methods the precision of EF was within the ~3-5% [46] needed when it is used as an index of LV function in clinical trials [47]. Lastly, we showed that the error in our measures of LV EDV and LVM was almost half the inter-observer (~10.6 mL, 12.0 g), and comparable to the intra-observer (~4.6 mL, 6.2 g) MAE reported in [31], but further investigations are required to assess the performance on more heterogeneous populations.

### B. Strain Measures

The application of myocardial strain to quantify abnormal deformation in disease requires accurate definition of normal ranges. However, previously reported normal ranges vary largely between modalities and techniques, particularly for radial ESS [4]. In this study we showed DeepStrain generated strain measures with narrow CI in healthy subjects from across three different datasets (Table 1). Although direct comparison with the literature is difficult due to differences in the datasets, overall our strain measures agreed with several reported results. Specifically, circumferential strain is in agreement with studies in healthy participants based on tagging (−16.6%, *n*=129) and speckle tracking echocardiography (−18%, *n*=265) datasets [48], [49], as well a recently proposed (−16.7% basal, *n*=386) tagging-based DL method [42]. Radial strain is in agreement with tagging-based (26.5%, *n*=129; 23.8% basal, *n*=386) studies [42], [48], but are lower than most reported values [4]. This is a result of smoothing regularization used during training to prevent overfitting. However, lowering the regularization without increasing the size of the training set would lead to increased EPE and wider CI. SR measures derived with DeepStrain were also in good agreement with previous tagging-based studies [48].

The CMAC dataset enabled us to compare our results to non-learning methods using a common dataset. We found that AEPE was lower with tagging-based techniques, reflecting the advantage of estimating cardiac motion from a grid of intrinsic tissues markers (i.e., grid tagging lines). Further, the tagging techniques also benefited from the fact that landmarks were placed at the center of the ventricle, whereas motion estimation from tagging data at the myocardial borders and in thin-walled regions of the LV is less accurate due to the spatial resolution of the tagging grid [4]. In addition, some of the tagging-MRI images did not enclosed the whole myocardium and some contained imaging artifacts, which resulted in strain artifacts towards the end of the cardiac cycle (Fig. 2c, black arrow). We found that MEVIS had the lowest AEPE, which could be a result of their image term (4) that penalizes phase shifts in the Fourier domain instead of intensity values, an approach that is less affected by desaturation (i.e., fading) of the tagging grid over time. The UPF approach also achieved a low AEPE using multimodal integration and 4D tracking to leverage the strengths of both modalities and improve temporal consistency [11]. Although this approach could in principle be recast as DL technique using recurrent neural networks [50], this would require a significant increase in the number of learnable parameters, therefore very large datasets would be needed to avoid overfitting.

Using MEVIS as the tagging reference standard, we found no significant differences in measures of radial and circumferential ESS (Fig. 2c). Temporally, we found significant differences in SR measures between the two techniques that could be due to drift errors in the MEVIS implementation, i.e., errors that accumulate in sequential implementations in which motion is estimated frame-by-frame [32]. Although we did not observe considerable improvements in AEPE compared to tagging- and cine-based methods, an important advantage of our approach is the reduced computational complexity (~13 sec in GPU) relative to the proposed MEVIS (1-2 h), IUCL (3-6 h), UPF (6 h) and INRIA (5 h) approaches [32]. Specifically, because once trained our network does not optimize for a specific test subject (i.e., it does not iterate on the cine-data to generate the desired output), centering, segmentation, and motion estimation for the entire cardiac cycle can be accomplished much faster (<2 min in CPU).

An additional advantage of non-iterative implementations is that we obtain deterministic results. Since this implies the exact same motion estimates are generated given the same input, we expect strain measures not to vary meaningfully if the anatomy and function remain fixed. Here we studied this property by evaluating the intra-scanner repeatability, an important aspect to consider when assessing the potential clinical utility of DeepStrain. Global measures of ESS showed excellent repeatability with narrow LoAs and with absolute RCs of less than 5% on average, and regional analyses also showed the average RC and aRC was less than 1% in more than half of the polar map segments, with the maximum difference being 7%. Finally, all SR measures showed good to excellent repeatability, except for SR which was moderate.

### C. Clinical Evaluation

DeepStrain could be applied in a wide range of clinical applications, e.g., automated extraction of imaging phenotypes from large-scale databases (e.g., UK biobank [51]). Such phenotypes include global and regional strain, which are important measures in the setting of existing dysfunction with preserved EF [3]. DeepStrain generated measures of global strain and SR over the entire cardiac cycle from a cohort of 100 subjects in <2 min (Fig. 5). These results showed that radial SRe was reduced in patients with HCM and ARV, despite having a normal or increased LV EF. Decreased SRe with normal EF is suggestive of subclinical LV diastolic dysfunction, which is in agreement with previous findings [52], [53]. Our results also showed DeepStrain-based maps could be used to characterize regional differences between groups (supplementary Fig. S6).

At an individual level (Fig. 6), we showed that in MI patients, polar segments with decreased circumferential strain matched myocardial regions with infarcted tissue. Further, we showed that the changes in regional strain due to MI can be both diffuse and focal. These abnormalities could be used to discriminate dysfunctional from functional myocardium [54], or as inputs for downstream classification algorithms [55]. More generally, DeepStrain could be used to extract interpretable features (e.g., strain and SR) for DL diagnostic algorithms [56], which would make understanding of the pathophysiological basis of classification more attainable [57].

### D. Study Limitations

A limitation of our study was the absence of important patient information (e.g., age), which would be needed for a more complete interpretation of our strain analysis results, for example to assess the differences in strain values found between the healthy subjects from the ACDC and CMAC datasets. However, using publicly available data enables the scientific community to more easily reproduce our findings, and compare our results to other techniques. Another limitation was the absence of longitudinal analyses, i.e., longitudinal strain was not reported because it is normally derived from long-axis cine-MRI data not available in the training dataset. The size of the datasets is another potential limitation. The number of patients used for training is much smaller than the number of trainable parameters, potentially resulting in some degree of overfitting. To correct this, the training set for motion estimation could be expanded by validating the proposed segmentation network on more heterogeneous populations. Also, while our repeatability results were promising despite testing in only a small number of subjects, repeatability in patient populations was not shown.

### E. Conclusion

We developed an end-to-end learning-based workflow for strain analysis that is fast, operator-independent, and leverages real-world data instead of making explicit assumptions about myocardial tissue properties or geometry. This approach enabled us to derive strain measures from new data without further training or parameter finetuning, and our measures were robust to image artifacts, repeatable, and comparable to those derive from dedicated tagging data. These technical and practical attributes position DeepStrain as an excellent candidate for use in routine clinical studies or data-driven research.

## Acknowledgment

We acknowledge the support of NVIDIA Corporation with the donation of the Titan X Pascal GPU used for this research. We also thank P. Jodoin (ACDC) and C. Tobon-Gomez (CMAC) for their assistance with the challenge datasets.

## References

[1] M. A. Konstam and F. M. Abboud, “Ejection Fraction: Misunderstood and Over-rated (Changing the Paradigm in Categorizing Heart Failure),” Circulation, vol. 135, no. 8, pp. 717–719, Feb. 2017.

[2] P. Claus, A. M. S. Omar, G. Pedrizzetti, P. P. Sengupta, and E. Nagel, “Tissue Tracking Technology for Assessing Cardiac Mechanics,” JACC: Cardiovascular Imaging, vol. 8, no. 12, pp. 1444–1460, Dec. 2015.

[3] O. A. Smiseth, H. Torp, A. Opdahl, K. H. Haugaa, and S. Urheim, “Myocardial strain imaging: how useful is it in clinical decision making?,” Eur Heart J, vol. 37, no. 15, pp. 1196–1207, Apr. 2016.

[4] M. S. Amzulescu, M. De Craene, H. Langet, A. Pasquet, D. Vancraeynest, A. C. Pouleur, J. L. Vanoverschelde, and B. L. Gerber, “Myocardial strain imaging: review of general principles, validation, and sources of discrepancies,” European Heart Journal - Cardiovascular Imaging, Mar. 2019.

[5] N. F. Osman, S. Sampath, E. Atalar, and J. L. Prince, “Imaging longitudinal cardiac strain on short-axis images using strain-encoded MRI,” Magn. Reson. Med., vol. 46, no. 2, pp. 324–334, Aug. 2001.

[6] D. Kim, W. D. Gilson, C. M. Kramer, and F. H. Epstein, “Myocardial Tissue Tracking with Two-dimensional Cine Displacement-encoded MR Imaging: Development and Initial Evaluation,” Radiology, vol. 230, no. 3, pp. 862–871, Mar. 2004.

[7] N. Risum, S. Ali, N. T. Olsen, C. Jons, M. G. Khouri, T. K. Lauridsen, Z. Samad, E. J. Velazquez, P. Sogaard, and J. Kisslo, “Variability of Global Left Ventricular Deformation Analysis Using Vendor Dependent and Independent Two-Dimensional Speckle-Tracking Software in Adults,” Journal of the American Society of Echocardiography, vol. 25, no. 11, pp. 1195–1203, Nov. 2012.

[8] A. Schuster, V.-C. Stahnke, C. Unterberg-Buchwald, J. T. Kowallick, P. Lamata, M. Steinmetz, S. Kutty, M. Fasshauer, W. Staab, J. M. Sohns, B. Bigalke, C. Ritter, G. Hasenfuß, P. Beerbaum, and J. Lotz, “Cardiovascular magnetic resonance feature-tracking assessment of myocardial mechanics: Intervendor agreement and considerations regarding reproducibility,” Clinical Radiology, vol. 70, no. 9, pp. 989–998, Sep. 2015.

[9] Wenzhe Shi, Xiahai Zhuang, Haiyan Wang, S. Duckett, D. V. N. Luong, C. Tobon-Gomez, KaiPin Tung, P. J. Edwards, K. S. Rhode, R. S. Razavi, S. Ourselin, and D. Rueckert, “A Comprehensive Cardiac Motion Estimation Framework Using Both Untagged and 3-D Tagged MR Images Based on Nonrigid Registration,” IEEE Trans. Med. Imaging, vol. 31, no. 6, pp. 1263–1275, Jun. 2012.

[10] G. Pedrizzetti, P. Claus, P. J. Kilner, and E. Nagel, “Principles of cardiovascular magnetic resonance feature tracking and echocardiographic speckle tracking for informed clinical use,” Journal of Cardiovascular Magnetic Resonance, vol. 18, no. 1, p. 51, Dec. 2016.

[11] M. De Craene, G. Piella, O. Camara, N. Duchateau, E. Silva, A. Doltra, J. D’hooge, J. Brugada, M. Sitges, and A. F. Frangi, “Temporal diffeomorphic free-form deformation: Application to motion and strain estimation from 3D echocardiography,” Medical Image Analysis, vol. 16, no. 2, pp. 427–450, Feb. 2012.

[12] T. Mansi, X. Pennec, M. Sermesant, H. Delingette, and N. Ayache, “iLogDemons: A Demons-Based Registration Algorithm for Tracking Incompressible Elastic Biological Tissues,” Int J Comput Vis, vol. 92, no. 1, pp. 92–111, Mar. 2011.

[13] R. Avazmohammadi, J. S. Soares, D. S. Li, T. Eperjesi, J. Pilla, R. C. Gorman, and M. S. Sacks, “On the in vivo systolic compressibility of left ventricular free wall myocardium in the normal and infarcted heart,” Journal of Biomechanics, vol. 107, p. 109767, Jun. 2020.

[14] V. Kumar, A. J. Ryu, A. Manduca, C. Rao, R. J. Gibbons, B. J. Gersh, K. Chandrasekaran, S. J. Asirvatham, P. A. Araoz, J. K. Oh, A. C. Egbe, A. Behfar, B. A. Borlaug, and N. S. Anavekar, “Cardiac MRI demonstrates compressibility in healthy myocardium but not in myocardium with reduced ejection fraction,” International Journal of Cardiology, vol. 322, pp. 278–283, Jan. 2021.

[15] B. Zhu, J. Z. Liu, B. R. Rosen, and M. S. Rosen, “Image reconstruction by domain transform manifold learning,” arXiv:1704.08841 [cs], Apr. 2017.

[16] P. Dong, B. Provencher, N. Basim, N. Piché, and M. Marsh, “Forget About Cleaning up Your Micrographs: Deep Learning Segmentation Is Robust to Image Artifacts,” Microsc Microanal, pp. 1–2, Jul. 2020.

[17] G. Simantiris and G. Tziritas, “Cardiac MRI Segmentation With a Dilated CNN Incorporating Domain-Specific Constraints,” IEEE J. Sel. Top. Signal Process., vol. 14, no. 6, pp. 1235–1243, Oct. 2020.

[18] F. Isensee, P. Jaeger, P. M. Full, I. Wolf, S. Engelhardt, and K. H. Maier-Hein, “Automatic Cardiac Disease Assessment on cine-MRI via Time-Series Segmentation and Domain Specific Features,” arXiv:1707.00587 [cs], vol. 10663, 2018.

[19] C. Zotti, Z. Luo, A. Lalande, and P.-M. Jodoin, “Convolutional Neural Network With Shape Prior Applied to Cardiac MRI Segmentation,” IEEE J. Biomed. Health Inform., vol. 23, no. 3, pp. 1119–1128, May 2019.

[20] M. Baldeon Calisto and S. K. Lai-Yuen, “AdaEn-Net: An ensemble of adaptive 2D–3D Fully Convolutional Networks for medical image segmentation,” Neural Networks, vol. 126, pp. 76–94, Jun. 2020.

[21] K. Hammouda, F. Khalifa, H. Abdeltawab, A. Elnakib, G. A. Giridharan, M. Zhu, C. K. Ng, S. Dassanayaka, M. Kong, H. E. Darwish, T. M. A. Mohamed, S. P. Jones, and A. El-Baz, “A New Framework for Performing Cardiac Strain Analysis from Cine MRI Imaging in Mice,” Sci Rep, vol. 10, no. 1, p. 7725, Dec. 2020.

[22] E. Puyol-Anton, B. Ruijsink, W. Bai, H. Langet, M. De Craene, J. A. Schnabel, P. Piro, A. P. King, and M. Sinclair, “Fully automated myocardial strain estimation from cine MRI using convolutional neural networks,” in 2018 IEEE 15th International Symposium on Biomedical Imaging (ISBI 2018), Washington, DC, 2018, pp. 1139–1143.

[23] C. Qin, W. Bai, J. Schlemper, S. E. Petersen, S. K. Piechnik, S. Neubauer, and D. Rueckert, “Joint Learning of Motion Estimation and Segmentation for Cardiac MR Image Sequences,” arXiv:1806.04066 [cs], Jun. 2018.

[24] M. Qiao, Y. Wang, Y. Guo, L. Huang, L. Xia, and Q. Tao, “Temporally coherent cardiac motion tracking from cine MRI: Traditional registration method and modern CNN method,” Med. Phys., vol. 47, no. 9, pp. 4189–4198, Sep. 2020.

[25] H. Yu, S. Sun, H. Yu, X. Chen, H. Shi, T. S. Huang, and T. Chen, “FOAL: Fast Online Adaptive Learning for Cardiac Motion Estimation,” in 2020 IEEE/CVF Conference on Computer Vision and Pattern Recognition (CVPR), Seattle, WA, USA, 2020, pp. 4312–4322.

[26] P. Chen, X. Chen, E. Z. Chen, H. Yu, T. Chen, and S. Sun, “Anatomy-Aware Cardiac Motion Estimation,” arXiv:2008.07579 [cs, eess], Aug. 2020.

[27] B. D. de Vos, F. F. Berendsen, M. A. Viergever, M. Staring, and I. Išgum, “End-to-End Unsupervised Deformable Image Registration with a Convolutional Neural Network,” arXiv:1704.06065 [cs], vol. 10553, pp. 204–212, 2017.

[28] M. A. Morales, D. Izquierdo-Garcia, I. Aganj, J. Kalpathy-Cramer, B. R. Rosen, and C. Catana, “Implementation and Validation of a Three-dimensional Cardiac Motion Estimation Network,” Radiology: Artificial Intelligence, vol. 1, no. 4, p. e180080, Jul. 2019.

[29] A. Østvik, E. Smistad, T. Espeland, E. A. R. Berg, and L. Lovstakken, “Automatic Myocardial Strain Imaging in Echocardiography Using Deep Learning,” in Deep Learning in Medical Image Analysis and Multimodal Learning for Clinical Decision Support, vol. 11045, D. Stoyanov, Z. Taylor, G. Carneiro, T. Syeda-Mahmood, A. Martel, L. Maier-Hein, J. M. R. S. Tavares, A. Bradley, J. P. Papa, V. Belagiannis, J. C. Nascimento, Z. Lu, S. Conjeti, M. Moradi, H. Greenspan, and A. Madabhushi, Eds. Cham: Springer International Publishing, 2018, pp. 309–316.

[30] J.-U. Voigt, G. Pedrizzetti, P. Lysyansky, T. H. Marwick, H. Houle, R. Baumann, S. Pedri, Y. Ito, Y. Abe, S. Metz, J. H. Song, J. Hamilton, P. P. Sengupta, T. J. Kolias, J. d’Hooge, G. P. Aurigemma, J. D. Thomas, and L. P. Badano, “Definitions for a common standard for 2D speckle tracking echocardiography: consensus document of the EACVI/ASE/Industry Task Force to standardize deformation imaging,” European Heart Journal - Cardiovascular Imaging, vol. 16, no. 1, pp. 1–11, Jan. 2015.

[31] O. Bernard, A. Lalande, C. Zotti, F. Cervenansky, X. Yang, P.-A. Heng, I. Cetin, K. Lekadir, O. Camara, M. A. Gonzalez Ballester, G. Sanroma, S. Napel, S. Petersen, G. Tziritas, E. Grinias, M. Khened, V. A. Kollerathu, G. Krishnamurthi, M.-M. Rohe, X. Pennec, M. Sermesant, F. Isensee, P. Jager, K. H. Maier-Hein, P. M. Full, I. Wolf, S. Engelhardt, C. F. Baumgartner, L. M. Koch, J. M. Wolterink, I. Isgum, Y. Jang, Y. Hong, J. Patravali, S. Jain, O. Humbert, and P.-M. Jodoin, “Deep Learning Techniques for Automatic MRI Cardiac Multi-Structures Segmentation and Diagnosis: Is the Problem Solved?,” IEEE Trans. Med. Imaging, vol. 37, no. 11, pp. 2514–2525, Nov. 2018.

[32] C. Tobon-Gomez, M. De Craene, K. McLeod, L. Tautz, W. Shi, A. Hennemuth, A. Prakosa, H. Wang, G. Carr-White, S. Kapetanakis, A. Lutz, V. Rasche, T. Schaeffter, C. Butakoff, O. Friman, T. Mansi, M. Sermesant, X. Zhuang, S. Ourselin, H.-O. Peitgen, X. Pennec, R. Razavi, D. Rueckert, A. F. Frangi, and K. S. Rhode, “Benchmarking framework for myocardial tracking and deformation algorithms: An open access database,” Medical Image Analysis, vol. 17, no. 6, pp. 632–648, Aug. 2013.

[33] American Heart Association Writing Group on Myocardial Segmentation and Registration for Cardiac Imaging:, M. D. Cerqueira, N. J. Weissman, V. Dilsizian, A. K. Jacobs, S. Kaul, W. K. Laskey, D. J. Pennell, J. A. Rumberger, T. Ryan, and M. S. Verani, “Standardized Myocardial Segmentation and Nomenclature for Tomographic Imaging of the Heart: A Statement for Healthcare Professionals From the Cardiac Imaging Committee of the Council on Clinical Cardiology of the American Heart Association,” Circulation, vol. 105, no. 4, pp. 539–542, Jan. 2002.

[34] M. Jaderberg, K. Simonyan, A. Zisserman, and K. Kavukcuoglu, “Spatial Transformer Networks,” arXiv:1506.02025 [cs], Jun. 2015.

[35] S. Ioffe and C. Szegedy, “Batch Normalization: Accelerating Deep Network Training by Reducing Internal Covariate Shift,” arXiv:1502.03167 [cs], Mar. 2015.

[36] B. Xu, N. Wang, T. Chen, and M. Li, “Empirical Evaluation of Rectified Activations in Convolutional Network,” arXiv:1505.00853 [cs, stat], Nov. 2015.

[37] K. He, X. Zhang, S. Ren, and J. Sun, “Deep Residual Learning for Image Recognition,” arXiv:1512.03385 [cs], Dec. 2015.

[38] W. P. Segars, G. Sturgeon, S. Mendonca, J. Grimes, and B. M. W. Tsui, “4D XCAT phantom for multimodality imaging research: 4D XCAT phantom for multimodality imaging research,” Medical Physics, vol. 37, no. 9, pp. 4902–4915, Aug. 2010.

[39] L. Wissmann, C. Santelli, W. P. Segars, and S. Kozerke, “MRXCAT: Realistic numerical phantoms for cardiovascular magnetic resonance,” Journal of Cardiovascular Magnetic Resonance, vol. 16, no. 1, Dec. 2014.

[40] L. Tautz, A. Hennemuth, and H.-O. Peitgen, “Motion Analysis with Quadrature Filter Based Registration of Tagged MRI Sequences,” in Statistical Atlases and Computational Models of the Heart. Imaging and Modelling Challenges, vol. 7085, O. Camara, E. Konukoglu, M. Pop, K. Rhode, M. Sermesant, and A. Young, Eds. Berlin, Heidelberg: Springer Berlin Heidelberg, 2012, pp. 78–87.

[41] K. McLeod, A. Prakosa, T. Mansi, M. Sermesant, and X. Pennec, “An Incompressible Log-Domain Demons Algorithm for Tracking Heart Tissue,” in Statistical Atlases and Computational Models of the Heart. Imaging and Modelling Challenges, vol. 7085, O. Camara, E. Konukoglu, M. Pop, K. Rhode, M. Sermesant, and A. Young, Eds. Berlin, Heidelberg: Springer Berlin Heidelberg, 2012, pp. 55–67.

[42] E. Ferdian, A. Suinesiaputra, K. Fung, N. Aung, E. Lukaschuk, A. Barutcu, E. Maclean, J. Paiva, S. K. Piechnik, S. Neubauer, S. E. Petersen, and A. A. Young, “Fully Automated Myocardial Strain Estimation from Cardiovascular MRI–tagged Images Using a Deep Learning Framework in the UK Biobank,” Radiology: Cardiothoracic Imaging, vol. 2, no. 1, p. e190032, Feb. 2020.

[43] R. Vallat, “Pingouin: statistics in Python,” JOSS, vol. 3, no. 31, p. 1026, Nov. 2018.

[44] N. Painchaud, Y. Skandarani, T. Judge, O. Bernard, A. Lalande, and P.-M. Jodoin, “Cardiac MRI Segmentation with Strong Anatomical Guarantees,” in Medical Image Computing and Computer Assisted Intervention – MICCAI 2019, vol. 11765, D. Shen, T. Liu, T. M. Peters, L. H. Staib, C. Essert, S. Zhou, P.-T. Yap, and A. Khan, Eds. Cham: Springer International Publishing, 2019, pp. 632–640.

[45] M. Khened, V. Alex, and G. Krishnamurthi, “Densely Connected Fully Convolutional Network for Short-Axis Cardiac Cine MR Image Segmentation and Heart Diagnosis Using Random Forest,” in Statistical Atlases and Computational Models of the Heart. ACDC and MMWHS Challenges, vol. 10663, M. Pop, M. Sermesant, P.-M. Jodoin, A. Lalande, X. Zhuang, G. Yang, A. Young, and O. Bernard, Eds. Cham: Springer International Publishing, 2018, pp. 140–151.

[46] J. A. San Román, J. Candell-Riera, R. Arnold, P. L. Sánchez, S. Aguadé-Bruix, J. Bermejo, A. Revilla, A. Villa, H. Cuéllar, C. Hernández, and F. Fernández-Avilés, “Quantitative Analysis of Left Ventricular Function as a Tool in Clinical Research. Theoretical Basis and Methodology,” Revista Española de Cardiología (English Edition), vol. 62, no. 5, pp. 535–551, May 2009.

[47] J. P. Kelly, R. J. Mentz, A. Mebazaa, A. A. Voors, J. Butler, L. Roessig,M. Fiuzat, F. Zannad, B. Pitt, C. M. O’Connor, and C. S. P. Lam, “Patient Selection in Heart Failure With Preserved Ejection Fraction Clinical Trials,” Journal of the American College of Cardiology, vol. 65, no. 16, pp. 1668–1682, Apr. 2015.

[48] B. A. Venkatesh, S. Donekal, K. Yoneyama, C. Wu, V. R. S. Fernandes, B. D. Rosen, M. L. Shehata, R. McClelland, D. A. Bluemke, and J. A. C. Lima, “Regional myocardial functional patterns: Quantitative tagged magnetic resonance imaging in an adult population free of cardiovascular risk factors: The multi-ethnic study of atherosclerosis (MESA): Reference Values of Strain From Tagged MRI,” J. Magn. Reson. Imaging, vol. 42, no. 1, pp. 153–159, Jul. 2015.

[49] D. Muraru, U. Cucchini, S. Mihăilă, M. H. Miglioranza, P. Aruta, G. Cavalli, A. Cecchetto, S. Padayattil-Josè, D. Peluso, S. Iliceto, and L. P. Badano, “Left Ventricular Myocardial Strain by Three-Dimensional Speckle-Tracking Echocardiography in Healthy Subjects: Reference Values and Analysis of Their Physiologic and Technical Determinants,” Journal of the American Society of Echocardiography, vol. 27, no. 8, pp. 858–871.e1, Aug. 2014.

[50] Z. Gan, J. Tang, and X. Yang, “Left Ventricle Motion Estimation Based on Unsupervised Recurrent Neural Network,” in 2019 IEEE International Conference on Bioinformatics and Biomedicine (BIBM), San Diego, CA, USA, 2019, pp. 2342–2349.

[51] A. Fry, T. J. Littlejohns, C. Sudlow, N. Doherty, L. Adamska, T. Sprosen, R. Collins, and N. E. Allen, “Comparison of Sociodemographic and Health-Related Characteristics of UK Biobank Participants With Those of the General Population,” American Journal of Epidemiology, vol. 186, no. 9, pp. 1026–1034, Nov. 2017.

[52] S. Chen, J. Yuan, S. Qiao, F. Duan, J. Zhang, and H. Wang, “Evaluation of Left Ventricular Diastolic Function by Global Strain Rate Imaging in Patients with Obstructive Hypertrophic Cardiomyopathy: A Simultaneous Speckle Tracking Echocardiography and Cardiac Catheterization Study,” Echocardiography, vol. 31, no. 5, pp. 615–622, May 2014.

[53] A. J. Marian and E. Braunwald, “Hypertrophic Cardiomyopathy: Genetics, Pathogenesis, Clinical Manifestations, Diagnosis, and Therapy,” Circ Res, vol. 121, no. 7, pp. 749–770, Sep. 2017.

[54] M. J. W. Götte, A. C. van Rossum, J. W. R. Twisk, J. P. A. Kuijer, J. T. Marcus, and C. A. Visser, “Quantification of regional contractile function after infarction: strain analysis superior to wall thickening analysis in discriminating infarct from remote myocardium,” Journal of the American College of Cardiology, vol. 37, no. 3, pp. 808–817, Mar. 2001.

[55] N. Zhang, G. Yang, Z. Gao, C. Xu, Y. Zhang, R. Shi, J. Keegan, L. Xu, H. Zhang, Z. Fan, and D. Firmin, “Deep Learning for Diagnosis of Chronic Myocardial Infarction on Nonenhanced Cardiac Cine MRI,” Radiology, vol. 291, no. 3, pp. 606–617, Jun. 2019.

[56] Q. Zheng, H. Delingette, and N. Ayache, “Explainable cardiac pathology classification on cine MRI with motion characterization by semi-supervised learning of apparent flow,” arXiv:1811.03433 [cs, stat], Mar. 2019.

[57] P. N. Kampaktsis and M. Vavuranakis, “Diastolic Function Evaluation,” JACC: Cardiovascular Imaging, vol. 13, no. 1, pp. 336–337, Jan. 2020.

