## Supplementary for "DeepStrain: A Deep Learning Workflow for the Automated Characterization of Cardiac Mechanics"

**Supplementary Sections and Figures**

**S1. Acquisition Protocol**

Subjects were recruited to undergo repeated scans. Standard cardiac MRI protocols were acquired on a 3T MRI system (Biograph mMR, Siemens Healthiness, Germany, Erlangen) in 4 healthy volunteers during two separate consecutive scan sessions on the same day. Written consent was obtaining from all volunteers with approval of the institutional review board (2018P002912) and in agreement with the Health Insurance Portability and Accountability Act (HIPAA) at the Massachusetts General Hospital. The major inclusion criteria were >21 years old, no history of cardiovascular diseases, and no MR contradictions. Each MRI protocol included standard localizer scans and a single slice cine acquisition in a long cardiac axis view, followed by a stack of short axis cines covering the heart from the base to the apex using a retrospectively ECG gated balanced steady-state gradient echo sequence (True FISP) with the following parameters; TR=38.61ms, TE=1.26ms, FA=28 degrees, FOV=277x340mm, matrix size=208x170, slice thickness=6mm, slice gap=6mm, FOV%=81.7%, bandwidth=925Hz with phase-encoding parallel imaging acceleration (PAT) of 2 and 25 cardiac frames over consecutive breath-holds. After each full protocol, the volunteers were asked to leave the scanner room before going back in for a second acquisition of exactly the same protocol.

**S2. Network Architectures**

Let **Ck** denote a Convolution-BatchNorm-PReLU layer with k filters. **CDk** denotes a Upsampling-Convolution-BatchNorm-PReLU layer with upsampling applied using nearest-neighbor interpolation of stride 2x2x2. Unless specified, all convolutions are 3x3x3 spatial filters applied with 1x1x1 stride. An encoding layer **Ek** consists of a Ck layer followed by a second Ck layer with stride 2x2x2. A third Ck layer follows but without BatchNorm-PReLU. The output is the residual connection made by element-wise addition of the second Ck layer before BatchNorm-PReLU are applied, and the third Ck layer. A decoding layer **Dk** consists of a Ck layer with (1x1x1)-sized filters followed by CDk and Ck layers. Thus, the 2x2x2 strided convolution in Ek downsamples by a factor of 2, whereas the upsampling operation with stride 2x2x2 in Dk upsamples by a factor of 2 With this notation, the encoder-decoder architecture common to all three networks consists of

**encoder**: E64-E128-E256-E512-E512-E512-E512. **decoder**: D512-D512-D512-D256-D128-D64.

After the last layer in the decoder, a final CDk layer without BatchNorm -PReLU and with (1x1x1)-sized filters is applied to map to the number of output channels (k=1 for VCN, 4 for CarSON, and 3 for CarMEN). Exceptions to the rules above are: (1) CarSON consists of two-dimensional operations, i.e., convolutions are 3x3 spatial filters applied with 1x1 stride. (2) For CarMEN all filters and strides in CarMEN were of size 1 along the third dimension since subjects had variable resolution along that dimension.

**S3. Optimization Experiments**

To verify the effects of smoothness and anatomical regularization and to find the optimal $\lambda$ values for $\mathcal{L}_{CarMEN}$, we generated 10 synthetic cine-MRI datasets with known ground-truth motion and tissue segmentations using the MR extended cardiac-torso (MRXCAT) [1], [2] as described in [3], each dataset consisting of two frames at end-diastole and end-systole. We trained CarMEN for 50 epochs using various regularization values and tested the models by evaluating the AEPE between ground-truth and predicted motion estimates. In agreement with non-learning literature, setting the smoothness weight to $\lambda_s = 0$ leads to highly irregular motion vectors (e.g., off by more than 90 degrees) relative to ground-truth (see supplementary figure S1). Setting the smoothness and anatomical weights to $\lambda_s = 0.1$, $\lambda_a = 0.1$ leads to smoother and better aligned vectors, but the magnitude of the vectors is decreased. Increasing the anatomical weight $\lambda_a = 0.5$ further improves the estimates by generating vectors with similar magnitude and orientation compared to the ground-truth. Qualitatively, increasing smoothness and anatomical regularization reduced AEPE (see supplementary figure S2 a, b). The effect of anatomical regularization on the strain estimates when we increased $\lambda_a$ from 0.1 to 0.5 was also verified, the latter having lower bias in both radial and circumferential ESS (see supplementary section S2 c, d). Thus, the optimal values $\lambda_i = 0.01$ , $\lambda_s = 0.5, \lambda_s = 0.1$ were used to train CarMEN.

### References


[1] W. P. Segars, G. Sturgeon, S. Mendonca, J. Grimes, and B. M. W. Tsui, "4D XCAT phantom for multimodality imaging research: 4D XCAT phantom for multimodality imaging research," *Medical Physics*, vol. 37, no. 9, pp. 4902–4915, Aug. 2010, doi: 10.1118/1.3480985.

[2] L. Wissmann, C. Santelli, W. P. Segars, and S. Kozerke, "MRXCAT: Realistic numerical phantoms for cardiovascular magnetic resonance," *Journal of Cardiovascular Magnetic Resonance*, vol. 16, no. 1, Dec. 2014, doi: 10.1186/s12968-014-0063-3.

[3] M. A. Morales, D. Izquierdo-Garcia, I. Aganj, J. Kalpathy-Cramer, B. R. Rosen, and C. Catana, "Implementation and Validation of a Three-dimensional Cardiac Motion Estimation Network," *Radiology: Artificial Intelligence*, vol. 1, no. 4, p. e180080, Jul. 2019, doi: 10.1148/ryai.2019180080.


SUPPLEMENTARY TABLE I
**INTRA-SCANNER REPEATABILITY OF REGIONAL STRAIN MEASURES.**

| | Region | circ ICC [95% CI] | circ LoA | rad ICC [95% CI] | rad LoA |
|---|---|---|---|---|---|
| 0 | Whole-Map Average | 0.831 [0.170-0.990] | [-1.5 0.6] | 0.996 [0.940-1.000] | [-0.2 1.2] |
| 1 | Basal Anterior | 0.975 [0.680-1.000] | [-1.0 0.7] | 0.970 [0.620-1.000] | [-0.8 1.8] |
| 2 | Basal Anteroseptal | 0.905 [0.130-0.990] | [-2.0 1.5] | 0.907 [0.140-0.990] | [-2.4 2.5] |
| 3 | Basal Inferoseptal | 0.883 [0.020-0.990] | [-2.8 1.0] | 0.984 [0.780-1.000] | [-0.7 1.8] |
| 4 | Basal Inferior | 0.844 [0.130-0.990] | [-2.2 0.7] | 0.958 [0.500-1.000] | [-3.6 3.2] |
| 5 | Basal Inferolateral | 0.487 [0.680-0.960] | [-2.0 0.6] | 0.988 [0.830-1.000] | [-1.6 1.9] |
| 6 | Basal Anterolateral | 0.926 [0.260-1.000] | [-1.7 1.2] | 0.976 [0.690-1.000] | [-1.4 3.7] |
| 7 | Mid-Anterior | 0.972 [0.640-1.000] | [-1.0 0.8] | 0.987 [0.820-1.000] | [-0.1 1.4] |
| 8 | Mid-Anteroseptal | 0.902 [0.110-0.990] | [-2.0 1.4] | 0.900 [0.100-0.990] | [-1.9 2.3] |
| 9 | Mid-Inferoseptal | 0.875 [0.020-0.990] | [-2.8 1.0] | 0.967 [0.580-1.000] | [-0.9 2.3] |
| 10 | Mid-Inferior | 0.853 [0.100-0.990] | [-2.2 0.7] | 0.955 [0.480-1.000] | [-3.5 3.6] |
| 11 | Mid-Inferolateral | 0.588 [0.600-0.970] | [-1.7 0.3] | 0.988 [0.830-1.000] | [-1.3 2.3] |
| 12 | Mid-Anterolateral | 0.944 [0.380-1.000] | [-1.5 1.0] | 0.986 [0.810-1.000] | [-0.5 3.4] |
| 13 | Apical-Anterior | 0.975 [0.680-1.000] | [-0.9 0.7] | 0.985 [0.790-1.000] | [-0.8 3.0] |
| 14 | Apical-Septal | 0.900 [0.110-0.990] | [-1.7 1.3] | 0.924 [0.240-0.990] | [-1.3 1.7] |
| 15 | Apical Inferior | 0.874 [0.020-0.990] | [-2.6 0.9] | 0.974 [0.660-1.000] | [-1.3 1.8] |
| 16 | Apical Lateral | 0.826 [0.190-0.990] | [-1.2 -0.2] | 0.977 [0.700-1.000] | [-2.6 2.9] |
| 17 | Apical | 0.852 [0.100-0.990] | [-1.5 0.5] | 0.994 [0.910-1.000] | [-0.6 1.2] |

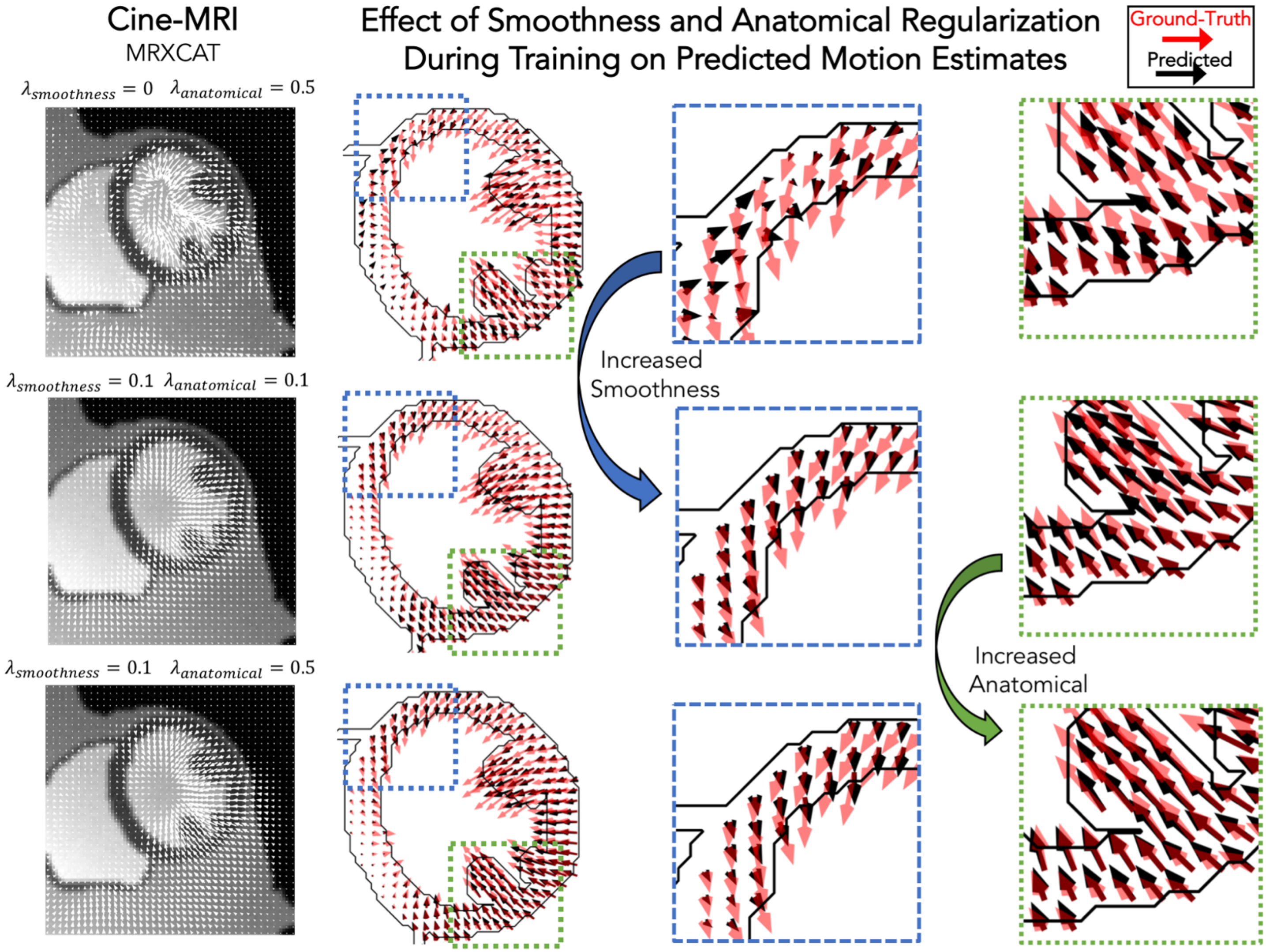


**Fig. S1 Qualitative effects of smoothing and anatomical regularization on the accuracy of motion estimates**. First row shows the predicted (black) motion estimates when the anatomical regularization is set to 0.5 and smoothing is set to 0. Relative to the ground-truth (red), these estimates are highly irregular. Increasing (third colum) the smoothness to 0.1 and setting anatomical to 0.1 improves the direction of the estimates, but the magnitude is reduced. This is corrected by increasing anatomical regularzation to 0.5 (fourth column).

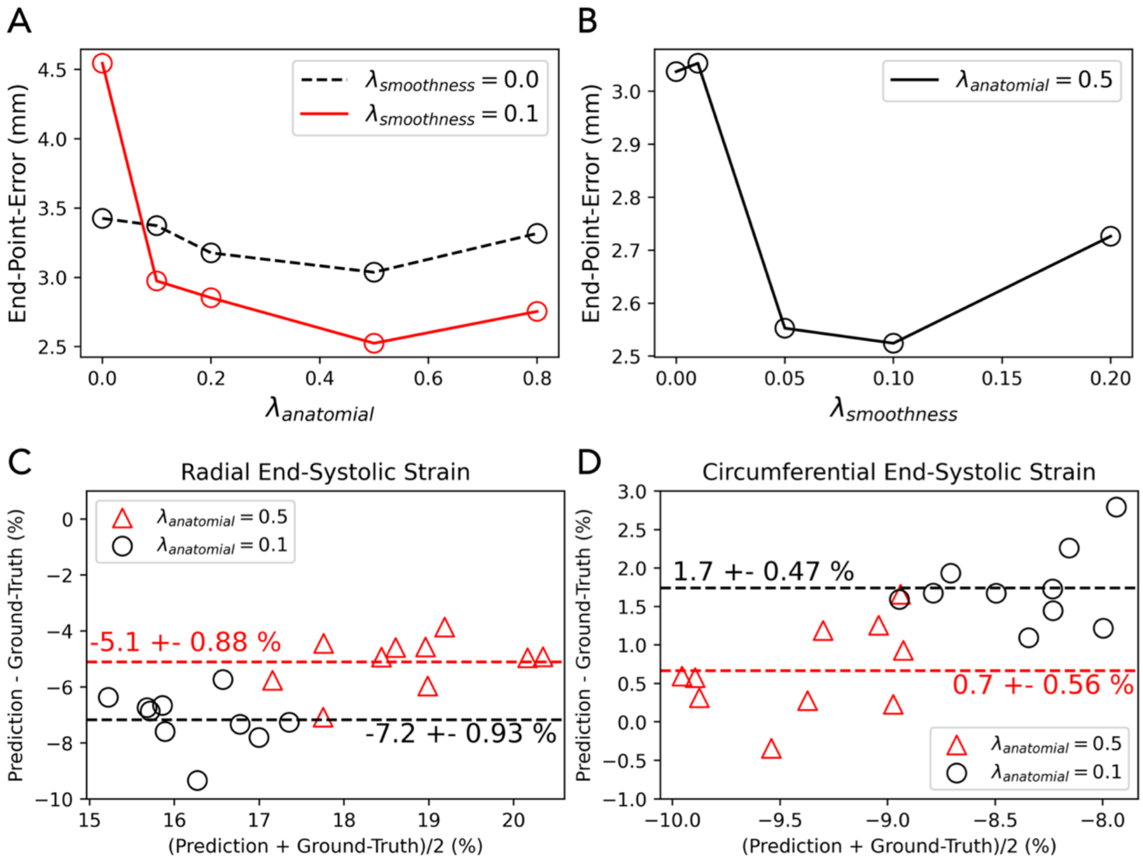


**Fig. S2 Quantitative effects of smoothing and anatomical regularization on the accuracy of motion and myocardial strain estimates.** Relative to the ground-truth motion vectors, the optimal (a) anatomical and (b) smoothing regularization parameters were found to be 0.5 and 0.1 based on the end-point-error. (c-d) Increasing the antomical regularization from 0.1 (black) to 0.5 (red) reduced biases in both the radial (c) and circumferential (d) end-systolic strain. Dotted line indicates mean difference between predicted and groud-truth strain values.

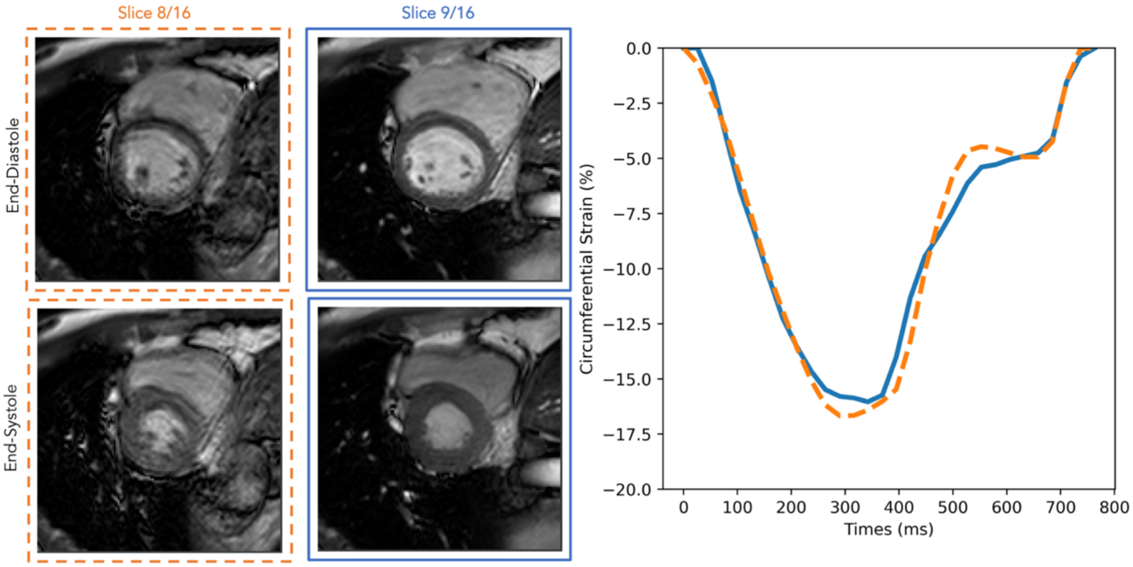


**Fig. S3 Effects of image artifacts on myocardial strain estimates.** In a healthy volunteer used to validate DeepStrain, imaging modality artifacts occurred during acquisition of slice 8 (left column), and were present at all time points (e.g., end-systole). Comparsion of the average strain, per-slice, against the adjacent slice without such artifacts (right column) showed little evidence that these artifacts affected the strain values derived with DeepStrain.

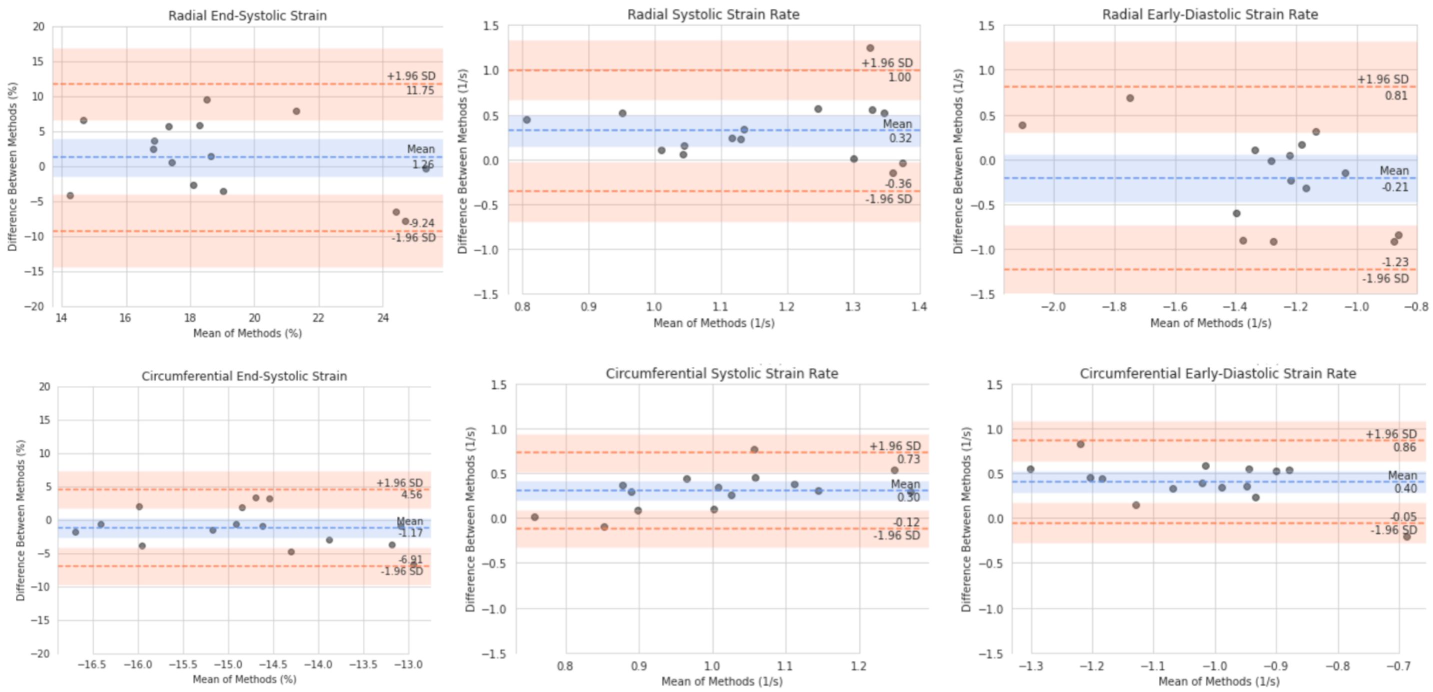


**Fig. S4 Validation of myocardial strain estimates using tagging-MRI reference method**. Bland-Altman plots of the strain measures obtained for the left ventricle at end-systole. The strain values obtained from DeepStrain were compared with those from the tagging reference standard. The first row shows the radial strain (first colum), systolic strain rate (second column), and early-diastolic strain rate (third column). The second row shows similar circumferential measures. Blue line denotes the mean difference; orange lines denote the 95% limits of agreement (mean ± 1.96 · standard devidation [SD]).

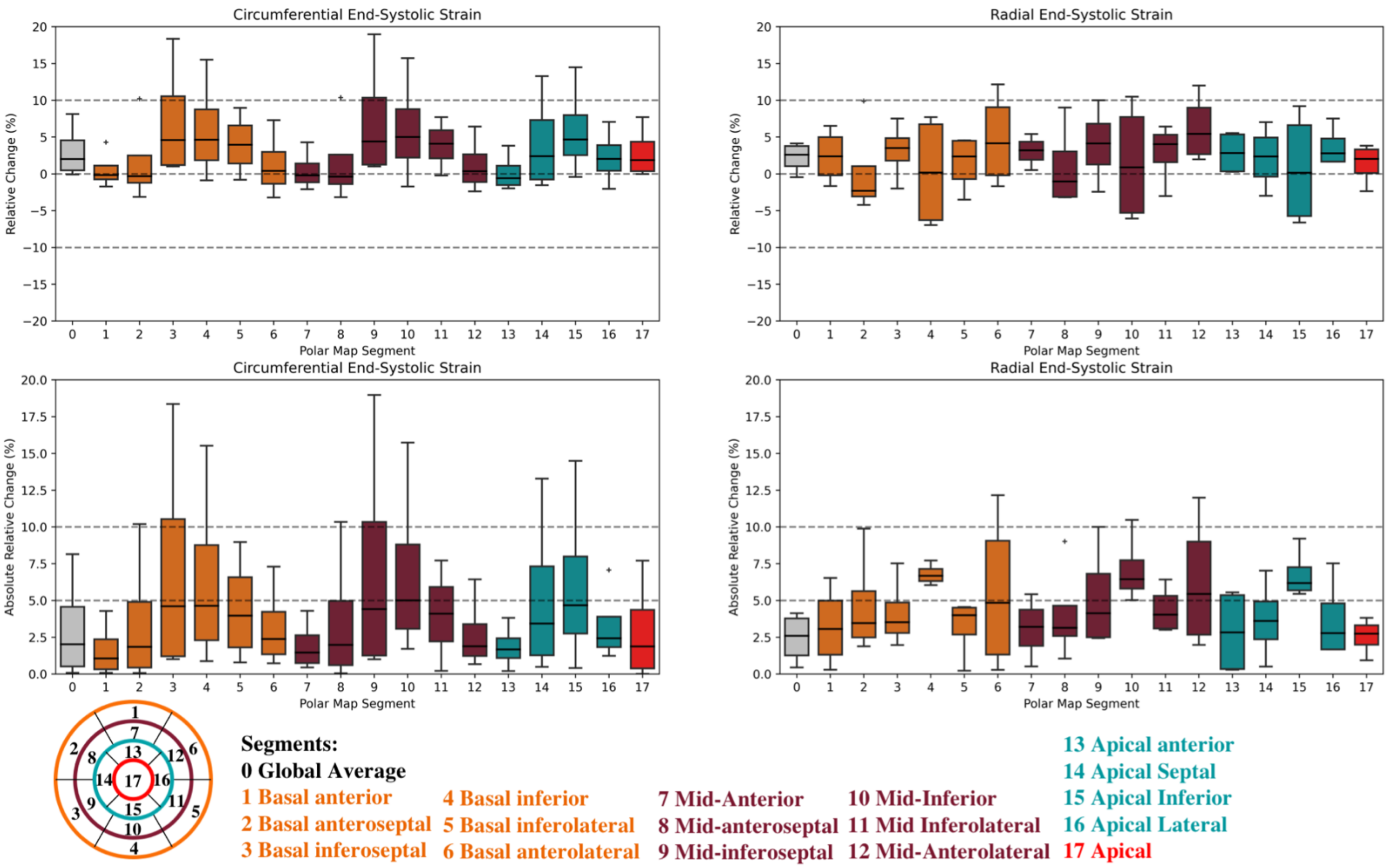


**Fig. S5 Segment-based intra-scanner repeatability of strain measures**. The first row shows the relative change (RC) of radial (first column) and circumferential (second column) end-systolic strain between acquisitions 1 and 2. The second row shows similar comparison via the absolute relative change (aRC). Each boxplot 1-17 corresponds to one of the American Heart Association polar map segments, 0 indicating the whole-map average.

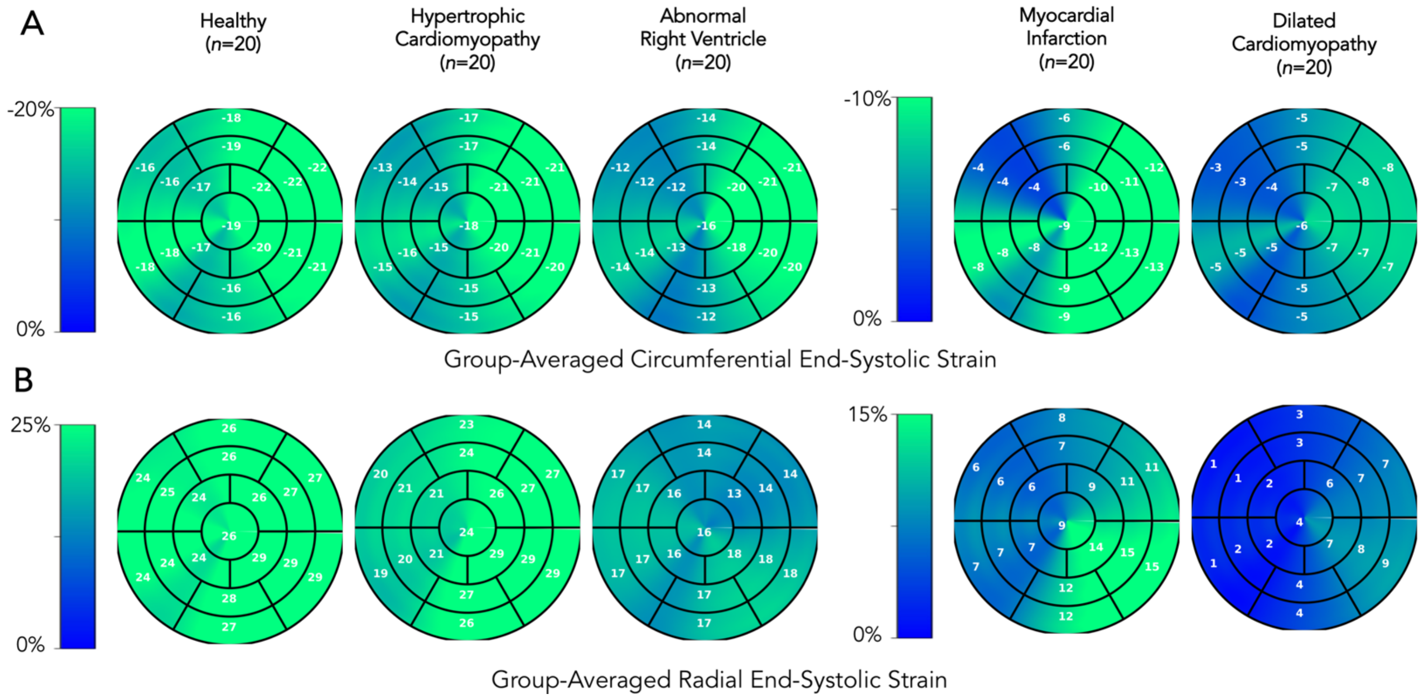


**Fig. S6 Group-wise comparison, a potential clinical application of regional strain**. Group-averaged regional strain values evaluated at end-systole for five groups shows progressive decline, starting with patients with hypertrophic cardiomyopathy (second column), and followed by patients with abnormal right ventricle, myocardial infarction, and dilated cardiomyopathy.
